# Specialisation of photosynthetic biochemistry in C_4_ chloroplasts potentiates distinct cell-specific functionality of different chloroplast-to-nucleus retrograde signalling pathways

**DOI:** 10.64898/2026.10.25.760861

**Authors:** Suyan Yee, Riley Furbank, Garima Bhatia, Su Yin Phua, Nikola Micic, Myat Thwe Naing, Nanna Bjarnholt, Nicholas Cox, Barry J. Pogson, Robert T. Furbank, Kai Xun Chan

## Abstract

Chloroplasts are key environmental sensors for plant cells. Upon exposure to abiotic stresses such as excess sunlight and heat, reactive oxygen species (ROS) in chloroplasts are sensed to trigger chloroplast-to-nucleus signalling pathways for acclimatory nuclear gene expression. There is extensive biochemical- and cell type-specialisation of chloroplasts, such as those in heat-resilient plants performing C_4_ photosynthesis, but how these specialisations intersect with chloroplast-to-nucleus signalling remains unclear. Here we show that specific chloroplast-to-nucleus signalling networks are functional in specialised leaf cell types of the C_4_ model species, *Setaria viridis*. RNA-sequencing, electron paramagnetic resonance (EPR) spectroscopy and metabolomics on cell type-enriched fractions, complemented with mutant analyses provided evidence for cell-specific enrichment of chloroplast-to-nucleus retrograde signalling pathways correlating with the photosynthetic specialisation in these cell types. Singlet oxygen (^1^O_2_)-responsive beta-cyclocitral (β-CC) /dihydroactinidiolide (DHA) signalling and hydrogen peroxide (H_2_O_2_)-responsive 3’-phosphoadenosine 5’-phosphate (PAP) signalling pathways were specific to mesophyll cells, in line with enrichment of photosystem II and linear electron flow to photosystem I, respectively in this cell type. In contrast, bundle sheath cells were enriched for the rapid, ROS-independent retrograde signalling pathway mediated by the Calvin cycle product, dihydroxyacetone phosphate (DHAP), consistent with the exclusive localisation of the Calvin cycle to C_4_ bundle sheath cells. Taken together, our results provide insights into how chloroplast- and cell type-specialisation intersect with stress signalling in plant cells and identify photosynthetic biochemistry as a key driver of cell-specific cellular signalling.

## INTRODUCTION

Adverse environmental conditions including heat, drought and excess sunlight induce the over-accumulation of reactive oxygen species (ROS), including singlet oxygen (^1^O_2_) and hydrogen peroxide (H_2_O_2_) in plant chloroplasts. The ROS over-accumulation alters the redox poise within plastids and causes oxidative damage to the photosynthetic apparatus, ultimately limiting photosynthesis and plant growth^1^. ROS over-production in chloroplasts can be sensed and communicated to the nucleus for activation of acclimation responses via several chloroplast-to-nucleus retrograde signalling pathways^2, 3^. At photosystem II (PSII), the over-production of ^1^O_2_ causes oxidative cleavage of beta-carotene to volatile beta-cyclocitral (β-CC) and dihydroactinidiolide (DHA), which signal via the cytosolic Methylene Blue Sensitivity 1 (MBS1) protein and the TGA2-Scarecrow-like 14 (SCL14) transcription factor complex to activate the expression of xenobiotic detoxification enzymes^4, 5^. A separate pathway for PSII-derived ^1^O_2_ sensing is mediated by the Executer 1 (EX1)/EX2 proteins which undergo ^1^O_2_-mediated tryptophan oxidation, leading to FtSH2-mediated proteolytic cleavage. The EX1/EX2 fragment(s) then function as a ^1^O_2_-responsive signal which modulate the expression of several genes involved in the regulation of cell death, including WRKY transcription factors^6, 7^. The β-CC/DHA and EX1/EX2 pathways are considered spatially separate as they respond to ^1^O_2_ produced at the PSII appressed thylakoids and grana margins, respectively^8, 9^.

The over-production of H_2_O_2_ due to photosystem I (PSI) can be sensed and signalled via multiple mechanisms^2, 10^. For the SAL-PAP pathway, overaccumulation of H_2_O_2_ leads to the oxidation of the cysteine residues in the adenosine phosphatase SAL enzyme^11^. H_2_O_2_-responsive disulfide bonding and modifications in SAL decreases its catabolic activity against its substrate, 3’-phosphoadenosine 5’-phosphate (PAP), a by-product of sulfotransferase (SOT) reactions.

The accumulated PAP functions as a retrograde signal by transport across the chloroplast membranes to the cytosol and nucleus. PAP inhibits the activity of nuclear exoribonucleases to actuate the expression of several ROS scavengers, such as Ascorbate Peroxidases (APXs) and the protein chaperone, Early Light-Induced Protein 2 (ELIP2)^12, 13^. Additionally, PSI-derived H_2_O_2_ can move directly to the nucleus via chloroplast-nuclear interactions or stromules^14, 15^. Alternatively, it can oxidize the 4Fe-4S iron-sulfur cluster of the 4-hydroxy-3-methylbut-2-enyl diphosphate synthase (HDS) enzyme that converts 2-*C*-methyl-D-erythritol 2,4-cyclodiphosphate (MEcPP) into (E)-4-hydroxy-3-methylbut-2-en-1-yl diphosphate (HMBPP) during isoprenoid biosynthesis^16^. This can lead to accumulation of MEcPP, which also functions as a retrograde signal to induce stress-responsive genes, such as *Hydroperoxide Lyase 1* (*HPL1*) and the signalling hub driven by the Calmodulin-binding transcription activator 3 (CAMTA3) transcription factor to activate general stress response genes, as well as genes associated with the unfolded protein response in the endoplasmic reticulum^17, 18^.

Chloroplasts also utilise several retrograde signalling pathways not directly linked to photosynthetic ROS^19, 20^. Upon illumination with high light intensity, the stimulation of photosynthetic light reactions leads to a rapid accumulation of the Calvin cycle intermediate, dihydroxyacetone phosphate (DHAP). DHAP is transported by the Triose Phosphate Translocator (TPT) into the cytosol, where it activates a MAP kinase signalling cascade to activate the expression and activity of several Apetela 2 (AP2)/Ethylene Response Factor (ERF) transcription factors^19^. This DHAP-AP2/ERF signalling cascade occurs rapidly and precedes ROS accumulation^21^. Additionally, chloroplast retrograde signalling pathways involved in chloroplast biogenesis, such as Genomes Uncoupled (GUN) 1, which regulates protein homeostasis^22, 23^ and GUN2 to GUN6 regulating heme production ^24, 25, 26^, have also been proposed to function in mature leaves and fully developed chloroplasts during high light stress^24, 27^. Recently, Moore and colleagues showed that high light stress induces NAD^+^/NADH-dependent, but ROS-independent, changes in translation of photosynthetic proteins in the cytosol^20^, presenting another mechanism for chloroplast signalling.

Largely, the aforementioned findings have focused on leaf-level signalling. Yet, there has been increasing indirect evidence suggesting that chloroplast retrograde signals play cell-specialised roles. For instance, constitutive PAP accumulation leads to lower ROS in leaf vascular bundles and is correlated with enhanced tolerance to high light stress^12^; whereas PAP induces higher ROS accumulation in guard cells, which triggers stomatal closure^28^. There are also significant differences in the proteomes of chloroplasts or plastids from mesophyll (M), bundle sheath (BS) and epidermal (E) pavement cells, with some retrograde signalling proteins being preferentially enriched in certain cell types^12, 29^. Thus, chloroplasts or plastids in some cell types, such as C_3_ BS and E pavement cells are proposed to not perform photosynthesis as their primary role and instead function as ‘sensory plastids’ to sense changes in the growth environment and influence plant responses via retrograde signalling^30^.

However, these observations of cell-specific signalling remain correlative and lack mechanistic detail. The extent and nature of structural and/or biochemical specialisations in chloroplasts and plastids across the different leaf cell types are still unclear in the C_3_ model plant *Arabidopsis*, where nearly all retrograde signalling studies to date have been performed. Several retrograde signals are derived from chloroplast metabolic pathways essential for photosynthesis (e.g. β-CC/DHA from carotenoids, MEcPP from isoprenoid biosynthesis, DHAP from the Calvin cycle). As such, it is difficult to envisage a mechanistic basis for cell-specialisation of retrograde signalling in plants performing C_3_ photosynthesis, such as *Arabidopsis*, where there has been no evidence of cell type-specialisation in photosynthetic light reactions or the Calvin cycle.

In contrast to C_3_ plants, the cell-type specialisation of chloroplast photosynthetic biochemistry has been extensively studied in plants performing C_4_ photosynthesis^31, 32^. C_4_ plants are characterised by a suite of integrated structural and biochemical adaptations, including Kranz-type leaf anatomy, specialised cell types, modified carbon metabolism and biochemistry, and distinct chloroplast architecture and function^33, 34, 35^. Together, these features enable the reactions of photosynthesis to be partitioned between M and BS cells, with each harbouring a distinct chloroplast type. Of the three C_4_ biochemical subtypes, the NADP-malic enzyme (NADP-ME) subtype is the most extensively studied, reflecting both its higher prevalence in agriculturally important crops, including maize, sorghum, and sugarcane, as well as its experimental simplicity, utility and tractability^36^. In particular, the extensive cell wall hardening in C_4_ BS cells has enabled high-purity mechanical separation of M and BS cells for study of these cell types individually^37^.

In the NADP-ME model plant *Setaria viridis*, M cells are specialised for photosynthetic light reactions, housing granal chloroplasts equipped with both PSI and PSII to support linear electron flow (LEF). In contrast, BS cells are specialised for the light-independent reactions of photosynthesis, housing agranal chloroplasts that are depleted in PSII and enriched in PSI, thereby favouring cyclic electron flow (CEF)^38, 39^. Calvin cycle enzymes are also spatially restricted to the BS chloroplasts in C_4_ species^32^. The dimorphic chloroplast architecture in C_4_ M and BS cells is accompanied by cell-specific differences in chloroplast number, organelle-to-organelle interactions, and cell wall modifications^40, 41^.

These extensive specialisations in chloroplast structure, distribution, and function between M and BS cells should necessitate corresponding alterations in chloroplast retrograde signalling pathways. However, apart from a link between increased chloroplast number in BS cells and chloroplast biogenic signalling^42, 43, 44^, virtually nothing is known about chloroplast retrograde signalling under abiotic stress in C_4_ plants. Here we utilise the C_4_ chloroplast dimorphism in M and BS cells of *S. viridis* to investigate how cell-type specialisation in photosynthetic biochemistry modulates active reprogramming of cellular signalling networks in response to environmental stimuli. We provide evidence for time-resolved and spatially distinct retrograde signalling pathways that are shaped by cell-specific photosynthetic specialisations in C_4_ M and BS cells.

## RESULTS

### Combined high light and heat treatment induces photosynthetic perturbations, ROS-responsive chloroplast signalling, and stress responses in *S. viridis*

Given that stress-responsive chloroplast signalling has not been explored in *S. viridis*, we first optimized a stress treatment regime suitable for interrogating short-term dynamic changes in chloroplast signalling in this species, on the scale of minutes-hours. Individual treatments of heat (H, 39°C) and high light (HL, 1200 μmol m^−2^ s^−1^) treatments alone had marginal effects on PSII quantum efficiency (YII) (**Fig S1A**), consistent with previous observations^45^. In contrast, combined high light and heat (HL+H, 1200 μmol m^−2^ s^−1^ at 39°C) treatment decreased YII by ∼50% and stimulated a corresponding increase in both non-regulated non-photochemical yield of PSII (YNO) and non-photochemical quenching (NPQ), indicative of photoinhibition and the engagement of photoprotection (**Fig S1A-C**). HL+H also induced a minor, but sustained decrease in PSI quantum efficiency (YI) (**Fig S1D**).

Consistent with these observations, HL alone did not significantly impact levels of ROS or the chloroplast signal PAP, whereas HL+H-treated plants exhibited significant ROS accumulation and ∼ 2-fold greater PAP accumulation than H-treated plants after 30 and 60 minutes of treatment (**Fig S2**). Together, these data suggest that compared to individual HL or H, combined HL+H treatment robustly impacts on photosynthetic electron transport to induce ROS-responsive chloroplast signalling in *S. viridis* leaves. Consequently, we utilised this HL+H treatment to investigate chloroplast signalling and gene expression in *S. viridis*.

### Cell type-resolved transcriptomes of *S. viridis* demonstrate specific and transient responses to high light and heat stress

To capture the dynamics of cell type-enriched stress responses in *S. viridis* to HL+H, we performed RNA sequencing (RNA-seq) on samples enriched in RNA from E, M or BS cells in addition to whole leaf samples (**Fig 1A**). Principal component analysis demonstrated clustering of biological replicates based on cell type and treatment, with these factors accounting for 13% and 75% of transcriptome variance, respectively (**Fig 1B**). Hierarchical clustering of samples based on Euclidian distance also highlighted a clear separation of samples between control and HL+H-stressed conditions, and the formation of finer sub-clusters corresponding to specific cell types (**Fig S3A**). We found the expected enrichment of cell type marker transcripts within each cell fraction^46^: *Carbonic Anhydrase* (*CA*), *phosphoenolpyruvate carboxylase (PEPC)*, and *NADP-dependent malate dehydrogenase* (*NADP-MDH*) were expressed preferentially within M samples, *Rubisco small subunit* (*RbcS*) and *phosphoribulokinase (PRK)* were enriched in BS samples (**Fig S3B**). The abundance of these photosynthesis-associated transcripts was minimal in E peel samples, consistent with their lack of green chloroplasts ^30^. The strong clustering of each cell type, as well as congruence with previous studies suggest our M, BS and E samples contained relatively little cross-contamination and are enriched for their respective cell types.

**Figure 1.**
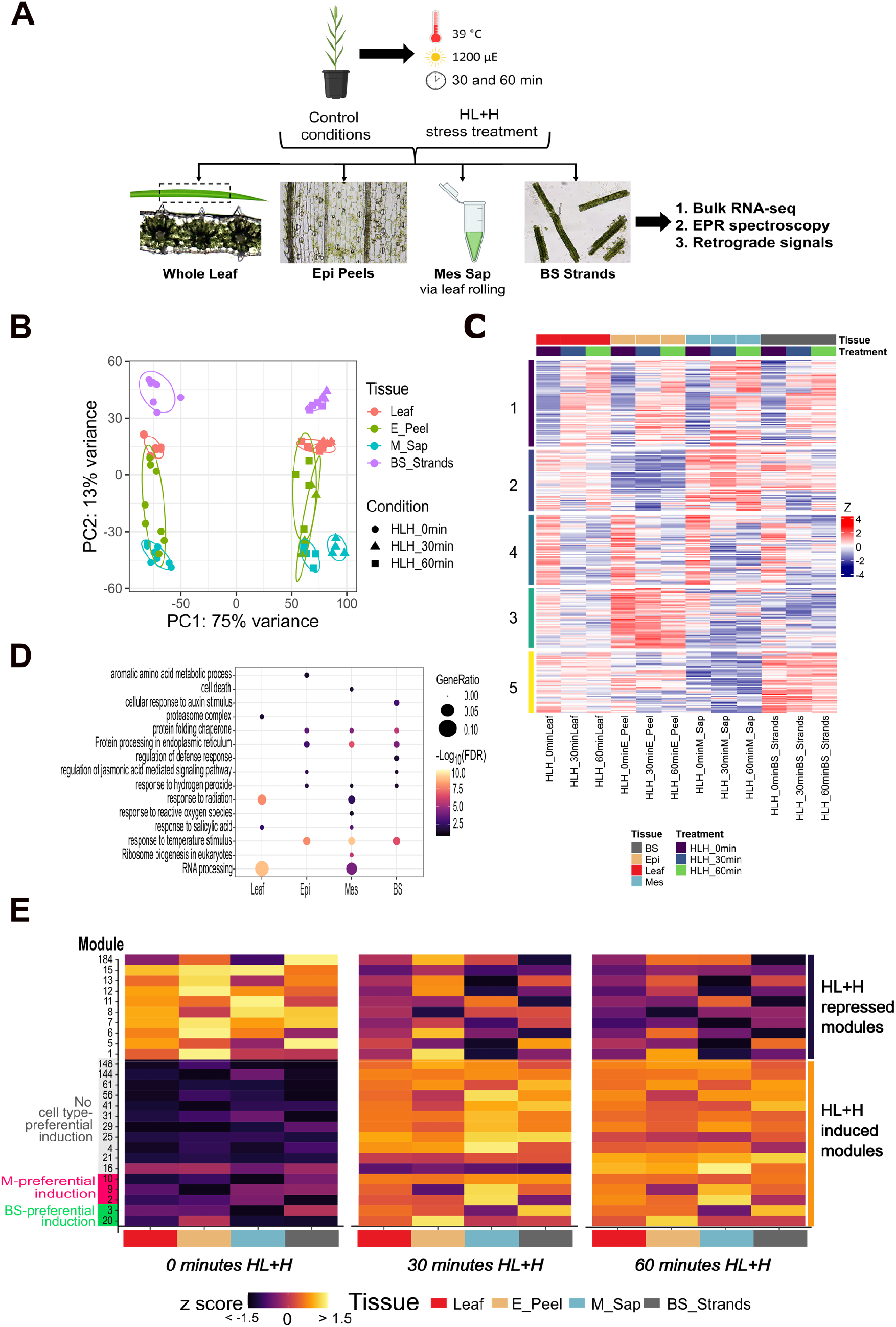
Cell type-enriched transcriptomics reveals varied and dynamic responses to high light and heat stress. **(A)** Overview of experimental workflow and sample preparation pipeline. Three-week-old *S. viridis* plants were harvested under control conditions, or treated with combined high light and heat for 30 or 60 minutes before harvesting whole leaves alongside cell type-enriched fractions for downstream RNA and metabolite analyses. **(B)** Principal component analysis of variance stabilized RNA seq counts. Centroid of each ellipse represents the mean ± 1 SD (approximately 68% of the data). *N* = 3-4 for each tissue:treatment combination. **(C)** Gene expression heatmap of all DEGs across the course of HL+H treatment. Colour scale represents the scaled Z-score of gene expression, with genes clustered into five groups via the K-means algorithm. All genes were significantly up-(adjusted p value < 0.05 & log_2_ fold change > |1|) or down-regulated (adjusted p value < 0.05 & log_2_ fold change < |1|) in response to either 30- or 60-minutes HL+H in at least one cell type. **(D)** Bubble plot of selected GO terms enriched under HL+H in different cell types. Gene ratio represents the percentage of DEGs present in this GO term divided by the total number of input genes. Colour scale represents the -Log_10_ transformed false discovery rate (FDR). **(E)** Annotated heatmap of gene co-expression modules organised by tissue and treatment type. Colour scale represents the mean Z-score of module gene expression in the given tissue:treatment combination.

To identify HL+H-responsive transcripts in each cell type, we next performed differential expression analysis using DESeq2 ^47^, classifying differentially expressed genes (DEGs) using a stringent cutoff of Log_2_ fold-change (LFC) > |1| and adjusted *P* value < 0.05. M cells had the highest number of unique DEGs with 2288, followed by the BS and E cells with 1435 and 1083 unique DEGs, respectively (**Fig S3C**). K-means clustering of all DEGs based on their LFC in response to HL+H treatment highlighted 5 general groups encompassing the core set of up- and down-regulated transcripts, in addition to smaller clusters defined by cell-specific patterns of differential expression (**Fig 1C, Supplementary Dataset S2**). Gene ontology (GO) enrichment analysis highlighted the over-representation of GO terms that are in common or unique to the different cell type enriched samples (**Fig 1D, Supplementary Dataset S3**).

To further interrogate cell-specific responses of the *S. viridis* transcriptome to HL+H, we constructed a Pearson correlation coefficient-based co-expression network encompassing all DEGs over the course of HL+H ^48^, where graph-based clustering grouped DEGs into 26 total co-expression modules (**Fig 1E**). The core set of HL+H-responsive DEGs common across all cell types represented as co-expression modules 4, 16, 21, 25, and 31 (**Fig S4, Supplementary Dataset S4-5**) formed a tightly clustered network. However, only clusters 16 and 21 had significantly enriched GO terms, which related to protein folding, RNA processing, and ribosome biogenesis (**Fig S4C**). To characterise broad differences in M and BS transcriptomic responses, we compared the HL+H-responsive LFC of DEGs in these cell types followed by GO analysis of cell-specific DEGs (**Fig S5, Supplementary Dataset S6**). After 30 minutes HL+H, the M-specific DEGs were enriched in terms related to the chloroplast, thylakoid membrane, and primary metabolism, while after 60 minutes of treatment, this shifted towards the proteasome and RNA modification-related terms. For BS-specific DEGs, GO terms were only significantly enriched after 30 minutes HL+H and included kinase activity and posttranslational modification, possibly suggesting that BS-specific responses are more transient (**Fig S5D**). BS responses to HL+H were specifically enriched in terms associated with hormone-mediated signalling and metabolism, while E responses were enriched in processes for aromatic amino acid biosynthesis (**Fig 1D**), consistent with the specialised functions of these cell types described by Hua *et al* ^49^ and Berkowitz *et al* ^50^. Interestingly, DEGs within M-enriched samples were uniquely enriched in the GO term “response to reactive oxygen species”, suggesting that biological processes related to oxidative stress and chloroplast signalling are not commonly upregulated across all cell types. Examination of the expression profiles of other co-expression modules highlighted several transcriptomic signatures that displayed cell type-enrichment (**Fig 1E**). These included modules 2, 9, and 10, all of which displayed strong M-preferential upregulation in response to HL+H, and modules 3 and 20, which displayed a transient BS-preferential upregulation after 30 minutes HL+H.

### β-cyclocitral-dependent chloroplast signalling mediates mesophyll-specific high light and heat responses

To identify if the cell-specific co-expression signatures related to further chloroplast signalling pathways, we analysed the M-preferential module 10 in greater detail (**Fig 2A**). GO analysis of module 10 transcripts revealed an enrichment of terms related to glutathione and UDP-glycosyltransferase activity (**Fig S6, Supplementary Dataset S7**). Strikingly, many of these terms were functionally related to xenobiotic detoxification processes, a key biochemical response driven by ^1^O_2_-dependent chloroplast signalling^4, 5^. Closer inspection of M-enriched DEGs revealed orthologues of *Scarecrow-like protein 14* (*SCL14*) and class II TGACG-binding (TGA) transcription factors, specifically *TGA6*, which together form a master regulatory complex activated upon β-CC treatment, in addition to transcripts encoding xenobiotic modifying enzymes and previously identified markers of β-CC signalling (**Fig 2B**). These transcripts were induced to a substantially lesser extent in BS strands (**Fig 2B**).

**Figure 2.**
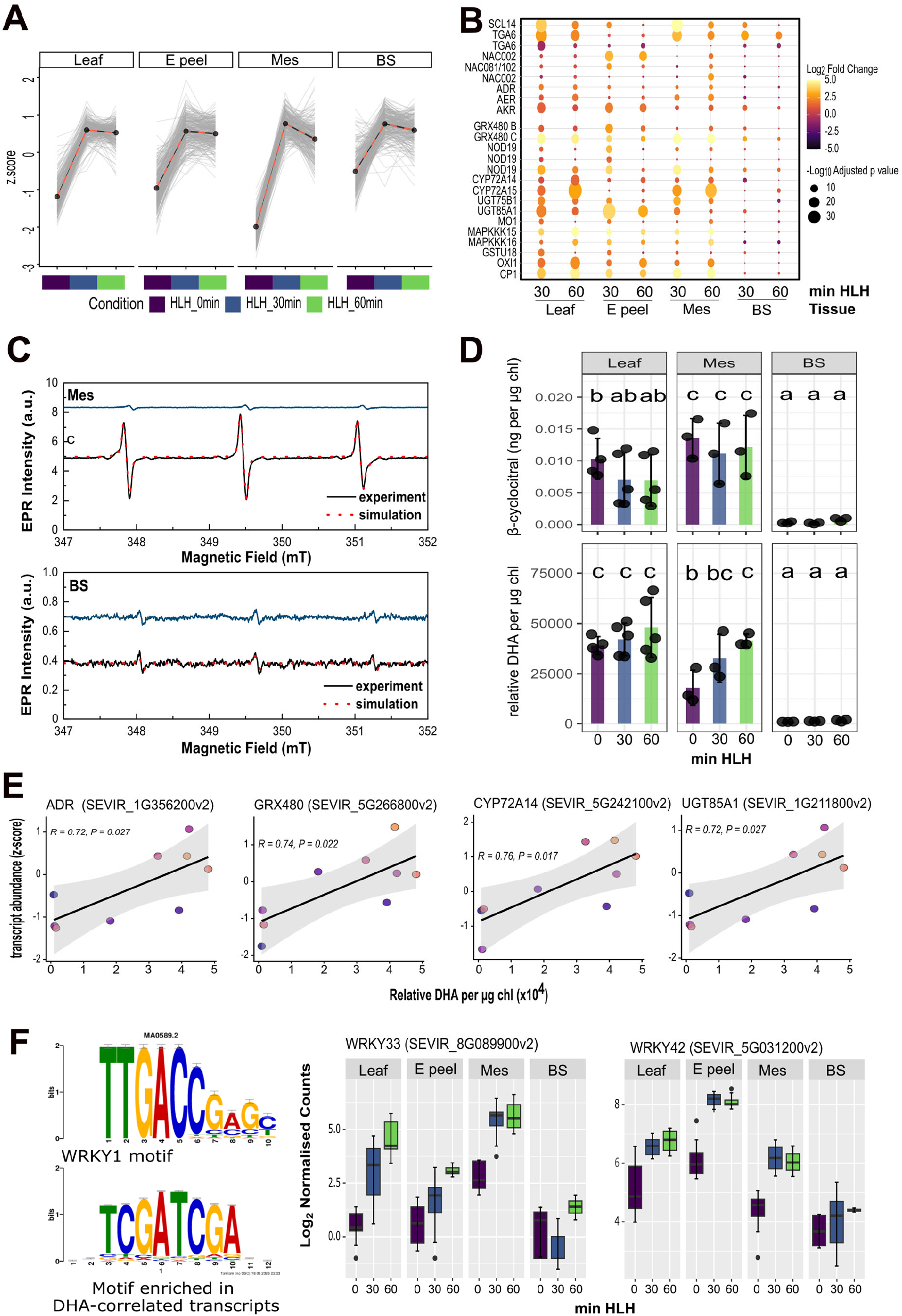
Mesophyll, but not bundle sheath cells, exhibit singlet oxygen-dependent chloroplast signalling under high light and heat stress. **(A)** Gene expression profile of co-expression module 10. Grey lines represent the expression patterns of each gene within the module (visualized as the genes z-score over the course of HL+H treatment [HLH]), and black lines show the mean expression calculated from all genes within the module. **(B)** Expression heatmap of genes related to singlet oxygen-dependent chloroplast signalling in response to high light and heat (HL+H). The colour scale represents the Log_2_-fold change in expression compared to 0 minutes HL+H treatment, and dot size represents the significance of this change as -Log_10_ of the adjusted *p*-value. **(C)** Representative EPR spectra of M or BS thylakoids in the presence of 100 µM 2,2,6,6-tetramethyl-4-piperidone (TEMPD) in the dark or after five minutes of high light treatment. Blue lines show residual EPR signal in the dark, solid black lines show signal after illumination, and dotted red points show the simulated spectra. **(D)** Absolute β-cyclocitral content (normalised to chlorophyll content) and relative dihydroactinidiolide content (normalised to chlorophyll content) of leaves, Mes sap and BS strands. Letters indicate statistically significant differences (Two-way ANOVA followed by Tukey’s post-hoc test, *p* < 0.05). **(E)** Correlation plots of DHA and xenobiotic detoxification marker transcript abundance. **(F)** Expression of two DHA-correlated WRKY genes and the WRKY motif (JASPAR ID MA0589.2) enriched in promoters of DHA-correlated transcripts.

We next investigated whether the M-enriched upregulation of transcripts related to xenobiotic detoxification and ^1^O_2_-responsive β-CC signalling is linked to the depletion of active PSII super-complexes, which produce ^1^O_2_, in the BS. To test this, we applied electron paramagnetic resonance (EPR) spectroscopy to the study of cell-specialised retrograde signalling for the first time. We utilised the spin-trap 2,2,6,6-tetramethyl-4-piperidone (TEMPD) to assess ^1^O_2_ production in isolated thylakoid membranes of M and BS cells. While both M and BS thylakoids exhibited low EPR signal in the dark, illumination with 1200 μmol m^−2^ s^−1^ white light induced a substantial increase in EPR signal in M thylakoids (**Fig 2C**). The EPR signal was consistent with the production of ^1^O_2_ as shown by the alignment to *in silico* simulated ^1^O_2_-responsive EPR spectra (**Fig 2C**). In contrast, only a small increase in EPR signal was observed in BS thylakoids.

We hypothesized that this cell-preferential accumulation of ^1^O_2_ could drive upregulation of ^1^O_2_- and β-CC/DHA-responsive genes in M sap through differential production of chloroplast signals like β-CC/DHA. We thus used headspace solid-phase microextraction coupled to gas chromatography and mass spectrometry (HS-SPME-GC-MS) to capture and quantify β-CC and DHA alongside other volatile organic compounds^51^. Mass-spectra of volatiles from leaf, M sap and BS strands were consistent with those of authentic standards, and similar amounts of a deuterium-labelled β-ionone volatile standard was recovered from spiked M and BS samples (**Fig S7**).

While β-CC was highly abundant in M sap and nearly undetectable in BS strands, neither β-CC nor its downstream metabolite beta-cyclocitric acid (β-CCA, which was not detected in any plant samples) significantly accumulated in response to HL+H, suggesting that β-CC may not be the predominant ^1^O_2_-responsive chloroplast signal induced under stress in *S. viridis* (**Fig 2D, Fig S7E**). However, DHA accumulated ∼2.5-fold after 60 minutes HL+H specifically in M sap and not in BS strands (**Fig 2D**). DHA accumulation was strongly correlated with expression of xenobiotic detoxification markers, as well as secondary metabolism and fungal response genes (**Fig 2E, Supplementary Dataset S8**). Among DHA-correlated transcripts, the most highly weighted regulators encoded transcription factors implicated in leaf senescence and ROS responses such as WRKYs 33 and 47^52, 53, 54^ (**Fig 2F**). Indeed, a motif closely resembling the WRKY-binding W-box was significantly enriched in the promoters of DHA-correlated transcripts (**Supplementary Dataset S9**). This motif occurred 8 times in the promoter of the DHA-correlated and ^1^O_2_ marker transcript GRX480 (SEVIR_5G266800v2), further suggesting a putative regulatory link between these processes and cell-specific ^1^O_2_ signalling. Taken together, these results suggest that BS cells have limited capacity for ^1^O_2_-responsive chloroplast signalling via β-CC or DHA, with xenobiotic detoxification and WRKY transcripts defining M-preferential responses to these signals.

### Mesophyll cells are hubs for PAP-dependent chloroplast signalling and reactive oxygen species detoxification

As PSI, the key site of photosynthetic H_2_O_2_ production, is present in both M and BS cells of *S. viridis*, we hypothesized that H_2_O_2_ and superoxide scavengers would thus be upregulated across both cells. However, analysis of specific transcripts encoding ROS scavengers highlighted a largely M- and E-preferential upregulation in response to HL+H (**Fig 3A**). These included *APX1*, the key peroxidase essential for cytosolic H_2_O_2_ scavenging, stromal APX, peroxiredoxin, and glutathione reductase. Transcripts encoding dehydroascorbate reductases (DHARs), monodehydroascorbate reductases (*MDAR*s), and markers of SAL-PAP signalling such as the *ELIP2*^12^ genes were also up-regulated to a greater extent in M cells than the BS (**Fig 3A**). This led us to hypothesize that cell-specific differences in PSI-derived ROS, such as H_2_O_2_ accumulation under HL+H may drive these transcriptomic responses. To test this, we monitored production of PSI-derived ROS using the spin trap 5,5-Dimethyl-1-Pyrroline-N-Oxide (DMPO) and EPR spectroscopy in isolated thylakoid membranes of M and BS cells. In response to illumination with 1200 μmol m^−2^ s^−1^ white light, M thylakoids exhibited a pronounced increase in signal primarily derived from the DMPO-OOH adduct as indicated by aN and aHβ hyperfine couplings of 1.42 and 1.15 respectively, consistent with substantial production of superoxide (**Fig 3B, Supplementary Dataset S10**). In contrast, only a minor increase was observed in BS thylakoids. These results suggest that under initial HL conditions, M cells may be more susceptible to PSI-derived ROS production, specifically superoxide, compared to BS cells.

**Figure 3.**
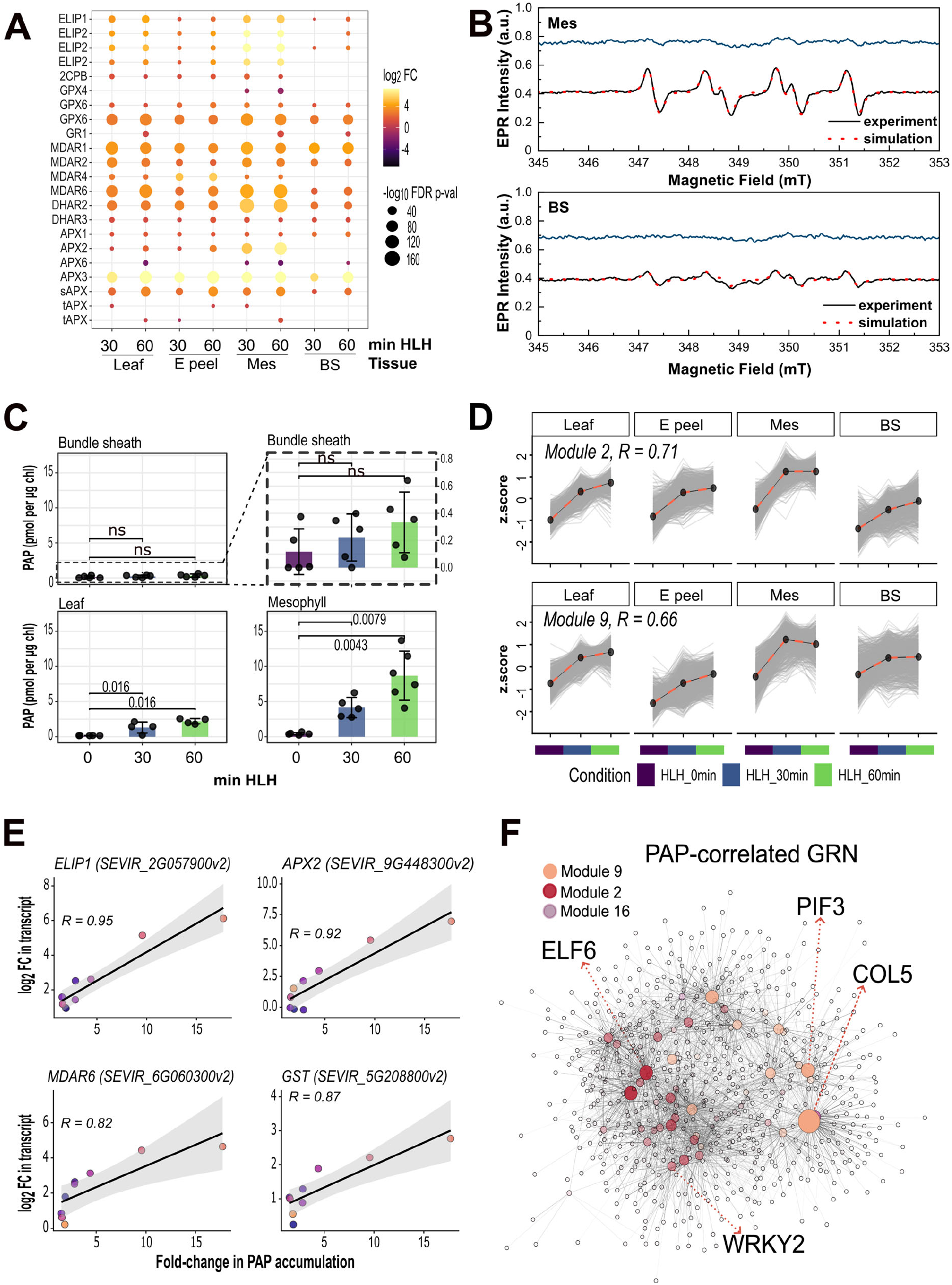
Mesophyll cells are hubs for PAP-dependent chloroplast signalling and reactive oxygen species detoxification. **(A)** Expression bubble plot of genes related to reactive oxygen species metabolism and PAP-dependent chloroplast signalling in response to high light and heat (HL+H). The colour scale represents the Log_2_-fold change in expression compared to 0 minutes HL+H treatment, and dot size shows the significance of this change as -Log_10_ of the adjusted *p*-value. **(B)** Representative EPR spectra of mesophyll and bundle sheath thylakoids in the presence of 50 mM DMPO in the dark and after 1-minute-high light treatment. Blue lines show residual EPR signal in the dark, solid black lines show signal after illumination, and dotted red points show the simulated spectra. **(C)** Absolute PAP levels (picomoles normalized to chlorophyll content) of leaves, mesophyll sap and BS strands after 0-, 30-, and 60-minutes HL+H treatment. Non-statistically significant differences between genotypes after a given duration of HL+H treatment are indicated by ns (Two-way ANOVA followed by Tukey’s post-hoc test, *p* < 0.05). **(D)** Gene expression profiles of the two significantly PAP-correlated co-expression modules and their associated Pearson correlation coefficient. Grey lines represent the expression patterns of each gene within the module (visualized as the genes z-score over the course of HL+H treatment), and black lines the mean expression calculated from all genes within the module. **(E)** Correlations between gene expression and PAP accumulation (represented by the Log_2_-fold change in expression and the fold-change in accumulation respectively) for four key PAP-associated transcripts. Pearson correlation coefficients are displayed, and the standard error of the fitted linear model shown using geom_smooth from ggplot2. **(F)** Gene regulatory sub-network of PAP-correlated transcripts. Colour of nodes represents their module membership within the co-expression network, and node size and transparency their regulatory weight in the gene regulatory network (GRN). Key hub regulators are annotated with *S. viridis* v2.0 transcript identifiers.

As SAL inactivation by chloroplast H_2_O_2_ leads to the accumulation of PAP^11^, we next asked whether this differential capacity for ROS production could drive M-specific PAP accumulation and associated transcriptomic remodelling. We thus quantified absolute PAP levels in cell type-enriched fractions and leaves in response to HL+H treatment. Treatment after 60 minutes HL+H resulted in an approximately 13-fold accumulation of PAP in *S. viridis* leaves (**Fig 3C**). Interestingly, while there was no significant PAP accumulation in the BS, a very large accumulation was observed in M cells, increasing approximately 10- and 20-fold after 30 and 60 minutes of HL+H respectively (**Fig 3C**), correlating with transcriptomic and EPR observations (**Fig 3A-B**). To identify additional targets of PAP signalling in *S. viridis*, we next integrated quantifications of PAP into the co-expression network and performed module-trait and gene-trait correlation analyses. This highlighted two key co-expression modules significantly correlated with PAP accumulation (modules 2 and 9), both of which exhibited M-preferential upregulation under HL+H and included established markers of PAP signalling, such as *APX2* and *ELIP* (**Fig 3D**). GO-enrichment analysis of PAP-correlated genes revealed terms associated with the chloroplast stroma and the photosynthetic membrane, mRNA processing, and splicing, consistent with findings that PAP alters expression of photosynthesis-associated nuclear genes and supports co-translational mRNA decay^55^ (**Fig S8, Supplementary Dataset S11-12**). To identify putative transcriptomic regulators of PAP signalling in M cells, we used Genie3 to generate a unidirectional gene regulatory network (GRN) based on our transcriptomic data and putative regulatory information, sub-setting this to include PAP correlated transcripts before visualising key regulatory interactions (**Fig 3F**). Highly weighted hub regulators in the network included orthologues of known stress-responsive WRKY transcription factors and several major photoperiod regulators, such as *Phytochrome-interacting factor 3* (*PIF3*), *Early Flowering 6* (*ELF6*), and *Constans-Like 5* (*COL5*) consistent with the established role of PAP in regulating the circadian system^56^.

As PAP accumulation is dependent on its production as a byproduct of secondary sulfur metabolism and catabolic activity of the SAL enzyme, we investigated whether transcripts implicated in these processes were differentially partitioned between M and BS cells. Overall, many transcripts related to primary sulfur assimilation, such as sulfate transporters (*SULTRs*), ATP sulfurylase (*ATPS*), and adenosine 5’-phosphosulfate reductase (*APR*) were enriched in BS strands, while the secondary sulfur metabolism enzymes facilitating PAP biosynthesis, such as APS kinases (*APKs*) and *SOTs* were more evenly distributed between cell types (**Fig S9**). We also identified two *SAL* genes in *S. viridis* (hereafter *SAL1* and *SAL2*), of which *SAL1* (SEVIR_7G291500v2) was expressed in both M sap and BS strands, while *SAL2* (SEVIR_7G291600v2) was enriched in BS strands and E peels. Both *SAL1* and *SAL2* are more highly expressed in BS compared to M. Differences in PAP catabolism via *SAL* and/or biosynthesis may thus be contributing factors to differential PAP accumulation in M and BS cells. Taken together, these data indicate that PAP-dependent chloroplast signalling and subsequent transcriptomic remodelling are largely M-, and to a lesser extent E-specific, responses to HL+H in *S. viridis*.

### The EXECUTER-, MEcPP- and GUN-dependent chloroplast signalling pathways may function in both mesophyll and bundle sheath cells

In addition to β-CC and DHA, ^1^O_2_ is also sensed via *EXECUTER 1 (EX1)* and *EX2*. *EX1* (SEVIR_9G214500v2) and one *EX2 ortholog* (*EX2A*, SEVIR_5G071200v2) transcripts were highly expressed in M cells and BS strands (**Fig S10A-C**). However, a second *EX2* ortholog (*EX2B,* SEVIR_3G102200v2) and the FtsH proteases necessary for EX1/2 cleavage^7, 57^ appear to be M-enriched (**Fig S10C-E**). Orthologues of many markers of *EX1*-dependent chloroplast signalling such as *WRKY*s *40* and *70* were also significantly upregulated in both M and BS cells (**Fig S10F**). Consequently, the M-specific up-regulation of ^1^O_2_-responsive transcripts appears to be specific to those mediated by β-CC signalling and not via *EX1/EX2*, with the latter potentially functioning in both cell types. Similarly, the ROS-responsive MEcPP retrograde signal may function in both M and BS cells as MEcPP-responsive genes were differentially regulated in both cell types, albeit with distinct subsets of MEcPP-responsive genes being induced in either M or BS cells (**Fig S11**).

The *Genomes Uncoupled (GUN)*-mediated retrograde signalling pathways, while predominantly known for their roles in chloroplast biogenesis, also participate in photooxidative stress responses, particularly in controlling the expression of *Light-Harvesting Chlorophyll a/b-Binding* (*LHCB*) transcripts^22, 24^. In our dataset, under control conditions nuclear-encoded *LHCB* transcripts were enriched in M-sap (**Fig S12A**) consistent with previous observations^46^. However, after 60 minutes HL+H both *LHCB1.5* and *LHCB3* were significantly repressed specifically in the M (**Fig S12A**). In contrast, some *LHCB* transcripts in the BS showed small, but statistically significant, increases in expression during HL+H treatment, such as *LHCB2.1* (**Fig S12A**). While neither *GUN2-6* genes nor the key *LHCB*-regulating transcription factor *Golden2-Like Kinase 1 (GLK1)* were differentially expressed under HL+H in M samples, *GUN1* was upregulated specifically in M sap while *GLK2* was repressed, coinciding with the M-specific repression of some *LHCBs* (**Fig S12B-C**). In contrast, in the BS, there is a small but significant repression of *GUN3* and *GUN4*, and small but significant upregulation in *GUN5* (**Fig S12B-C**). Genes co-expressed alongside *GUN1* exhibited similar HL+H-dependent upregulation in M sap, and to a lesser extent in E peels, and were enriched in GO terms related to light and UV responses, chloroplast organisation, and circadian rhythms (**Fig S12D-E, Supplementary Dataset S13**). Collectively, the transcriptional data suggest that *GUN*-mediated signalling may function in both M and BS cells, although the nature of their action in both cell types remained unclear.

### The DHAP-AP2/ERF rapid chloroplast signalling pathway underpins bundle sheath-specific responses to high light and heat

We next sought to characterize the transcriptomic responses and potential chloroplast signalling pathways that drive BS-specific changes under HL+H by analysing the co-expression module most closely correlated with BS-specific transcriptome changes in module 3 (**Fig 1E**). The genes in module 3 showed a significantly greater HL+H-dependent upregulation in BS and leaf tissue compared to M, with expression peaking in the BS after 30 minutes HL+H (**Fig 4A**). To interrogate potential transcriptional regulators driving these BS-specific expression profiles, we subset the previously generated GRN and visualized key regulatory interactions between hub transcription factors and module 3 genes. Transcripts belonging to module 3 formed a cluster within the network distinct from PAP-correlated and M-upregulated transcripts (modules 2 and 9), suggesting that unique regulatory interactions drive BS-specific transcriptomic responses to HL+H (**Fig 4B**). Within the module 3 regulatory network, the ethylene-responsive transcription factor (ERF) *RAP2.4* was the most highly weighted hub with 719 connections. Interestingly, several additional AP2/ERF-domain transcription factors were highly interconnected within the GRN and co-expressed alongside known stress-responsive transcripts such as calmodulins, NADPH-oxidases, and WRKY transcription factors. In Arabidopsis, several of these APR2/ERF-domain transcription factors are known to be activated by the rapid high light-responsive chloroplast retrograde signal, DHAP ^19^.

**Figure 4.**
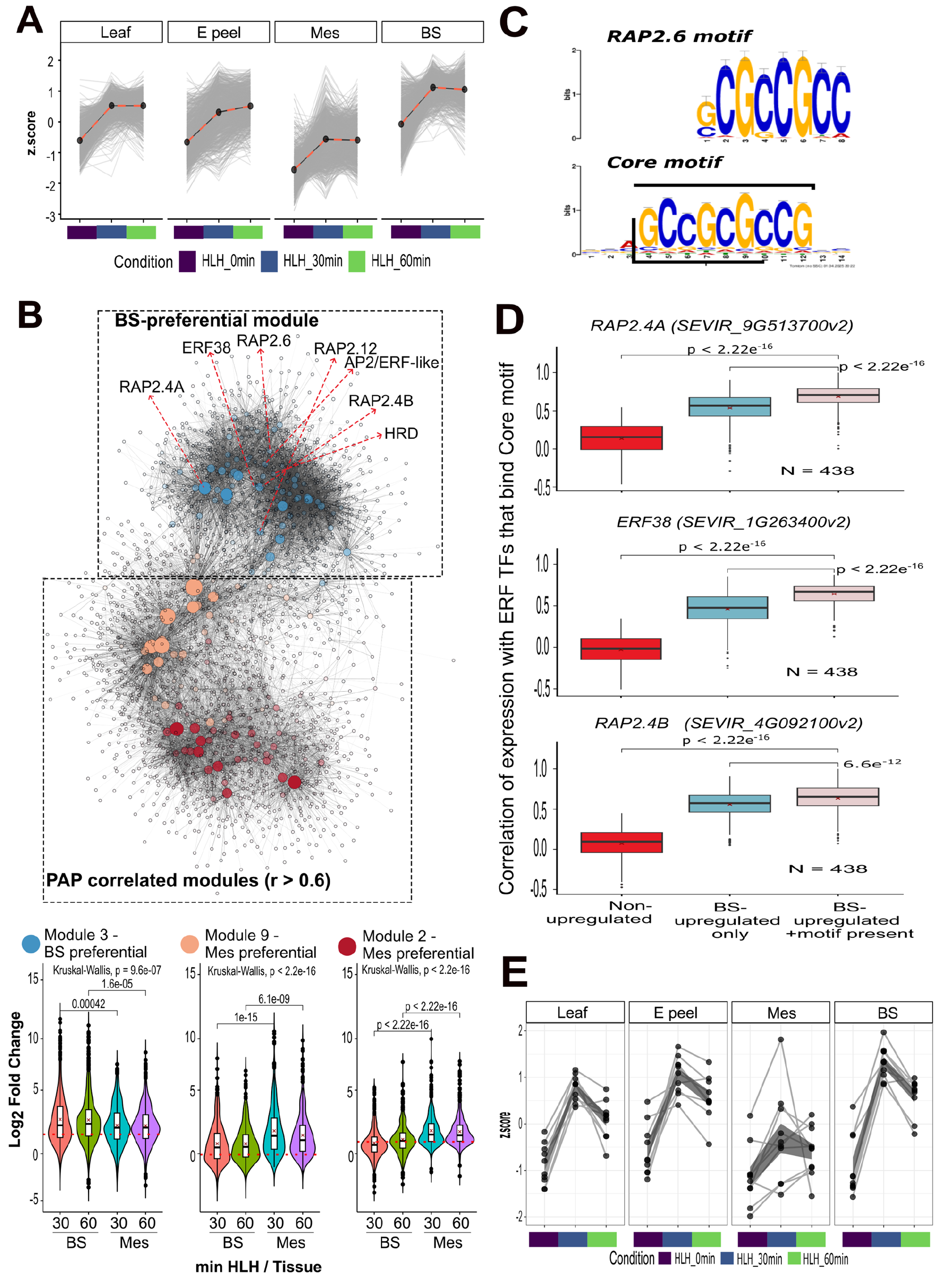
Bundle sheath responses to high light and heat are characterized by a network of AP2/ERF transcription factors and rapid, DHAP-dependent chloroplast signalling. **(A)** Gene expression profile of co-expression module 3. Grey lines represent the expression patterns of each gene within the module (visualized as the genes z-score over the course of HL+H treatment), and dotted lines the mean expression calculated from all genes within the module. **(B)** Gene regulatory sub-network consisting of genes belonging to the PAP-correlated co-expression modules 2 and 9, and the distinct BS-preferential module 3, the stress-responsive gene expression values of which are shown in the lower violin plots. Red dashed line in violin plots marks a log_2_ fold change of |1|, the cutoff for differential expression. Colour of nodes represents their module membership within the co-expression network, and node size their regulatory weight in the GRN. Key AP2/ERF family hub regulators are annotated with red arrows. **(C)** Graphical representation of workflow for transcription factor binding motif analysis, and the most enriched motif identified in module 3 transcripts aligned to that of RAP2.6 (JASPAR entry MA1052.1). **(D)** Pairwise correlations between AP2/ERF transcription factors and their putative targets within module 3. The correlation between AP2/ERF and downstream target gene expression was compared between 3 groups; transcripts within module 3 that contained the binding motif in **C** in their upstream promoter regions, module 3 transcripts that did not contain this motif in their promoter, and randomly selected transcripts from outside of module 3 (*N* = 438 for each group). Statistical significance was determined via Student’s two-tailed *t*-test and corrected for multiple comparisons using FDR. **(E)** Expression profiles (as scaled Z-scores) of APATELA2/Ethylene response transcription factors (AP2/ERF TFs) potentially involved in DHAP-dependent chloroplast signalling. Each grey line follows the expression of a distinct AP2/ERF transcript over the course of HL+H stress, with black points indicating the mean Z-score at each stress timepoint. Dark grey bands show the standard error across all genes at each time point. All genes were significantly differentially expressed in at least one cell type:treatment combination.

To investigate a potential regulatory link between DHAP-activated AP2/ERF transcription factors and BS-specific responses to HL+H, we extracted promoter sequences for all transcripts present in co-expression module 3 and performed de-novo motif enrichment analysis. The most significantly enriched motif in these sequences is displayed in **Fig 4C**, which contained the GCC-box commonly associated with ERF transcription factors, such as *RAP2.6* (**Supplementary Dataset S14**)^58^. Of the 921 genes within module 3, 438 (48% in total) contained this motif in their promoters. To test this putative regulatory link between AP2/ERF transcription factors and BS-upregulated transcripts, we examined correlations between the transcription factors associated with this motif, specifically *ERF110*, *RAP2.4*, and *ERF38*, and the module transcripts containing the GCC-box motif (**Fig 4D, Fig S13**). While it was expected that expression of these transcription factors would be strongly correlated with other transcripts preferentially upregulated in the BS, we hypothesized that correlations would be significantly greater in transcripts that also contained the associated GCC-box motif in their promoters (N = 438). As a negative control, we also tested a group of 438 randomly selected transcripts outside co-expression module 3 and not co-expressed alongside these transcription factors.

Strikingly, for five of six transcription factors tested, we observed significantly higher correlations between the transcription factors and transcripts that were both co-expressed and contained the motif compared to the other two groups (**Fig 4D, Fig S13**). This suggests a potential regulatory role of stress-inducible AP2/ERF transcription factors in mediating BS-specific stress responses. Indeed, many of these AP2/ERFs were transiently up regulated after 30 minutes HL+H, before decreasing in expression to near control levels by 60 minutes, a pattern that was consistent within the BS, E, and leaf, but not in M cells (**Fig 4E**). Among these were orthologues of *Redox-Responsive Transcription Factor 1* (*RRTF1*), *ERF38*, and *RAP2.4*, all of which are transiently upregulated by HL in *A. thaliana*^19, 59^. Furthermore, co-expression modules 3 and 20 which exhibited BS-preferential patterns of upregulation contained 2.1 and 4.5 times as many AP2/ERF transcripts as would be expected by random chance (Fisher exact test, *P* < 0.05). Interestingly, module 3 DEGs were enriched in GO terms related to oxoacid and carboxylic acid metabolism, while transcripts containing the GCC-box motif were specifically enriched for carbohydrate transport and secondary metabolites, consistent with DHAP being derived from the Calvin cycle and suggesting that this AP2/ERF network may regulate BS-specific changes in primary carbon metabolism and antioxidant biosynthesis (**Fig S14**, **Supplementary Dataset S15-16**). Taken together, these data suggest that a tight network of DHAP-responsive AP2/ERF transcription factors drive rapid BS-specific responses to HL+H.

### *MBS1* mediates mesophyll-specific responses to high light and heat

Having observed clear differential partitioning of chloroplast signals and their downstream transcriptional signatures between M and BS cells, we sought to genetically test the contribution of one of these putative cell-specific signalling pathways to abiotic stress responses. We targeted the MBS1 protein, which is a key signal transducer downstream of both DHA and β-CC, mediating the stress-responsive expression of several ^1^O_2_-responsive genes in *A. thaliana*^60, 61^. Surprisingly, despite the M-specific DHA accumulation and ^1^O_2_-responsive gene expression (**Fig 2**), *MBS1* expression was not significantly M-enriched (**Fig S15**).

We generated gene-edited *mbs1* mutants via CRISPR-Cas9 (**Fig S16A**). The gene-edited *mbs1* mutants showed frame shift mutations which altered the protein sequence, resulting in loss of stably functional MBS1 protein during HL+H treatment (**Fig S16B-C**). Whole-leaf PSII chlorophyll fluorescence measurements, which serve as a proxy for M photosynthetic performance due to PSII enrichment in this cell type, were then performed on WT and two *mbs1* null mutants (hereafter *mbs1* A and B). PSII quantum yield (YII) of both *mbs1* null mutants was comparable to WT under control conditions (**Fig 5A**). However, after 60 minutes HL+H treatment *mbs1 A* plants had significantly lower (YII) values. Strikingly, after 60 minutes of HL+H both *mbs1* mutants also had significantly lower yields of NPQ even under lower irradiance, which was reflected in higher YNO values indicating sustained photodamage (**Fig 5B-D**). Furthermore, *mbs1* plants had hampered induction of M-specific DHA-responsive marker genes, such as *Calcium-binding Protein 1 (CP1)* and the glycoside hydrolase *BGLU11* in response to HL+H, and constitutive induction of *Senescence Associated Gene 21 (SAG21)*, indicating disrupted ^1^O_2_-dependent signalling under both control and stress conditions (**Fig 5E**). Together, these results reinforce the importance of cell type-specific chloroplast signalling pathways, such as^1^O_2_-responsive DHA signalling, for *S. viridis* acclimation to HL+H.

**Figure 5.**
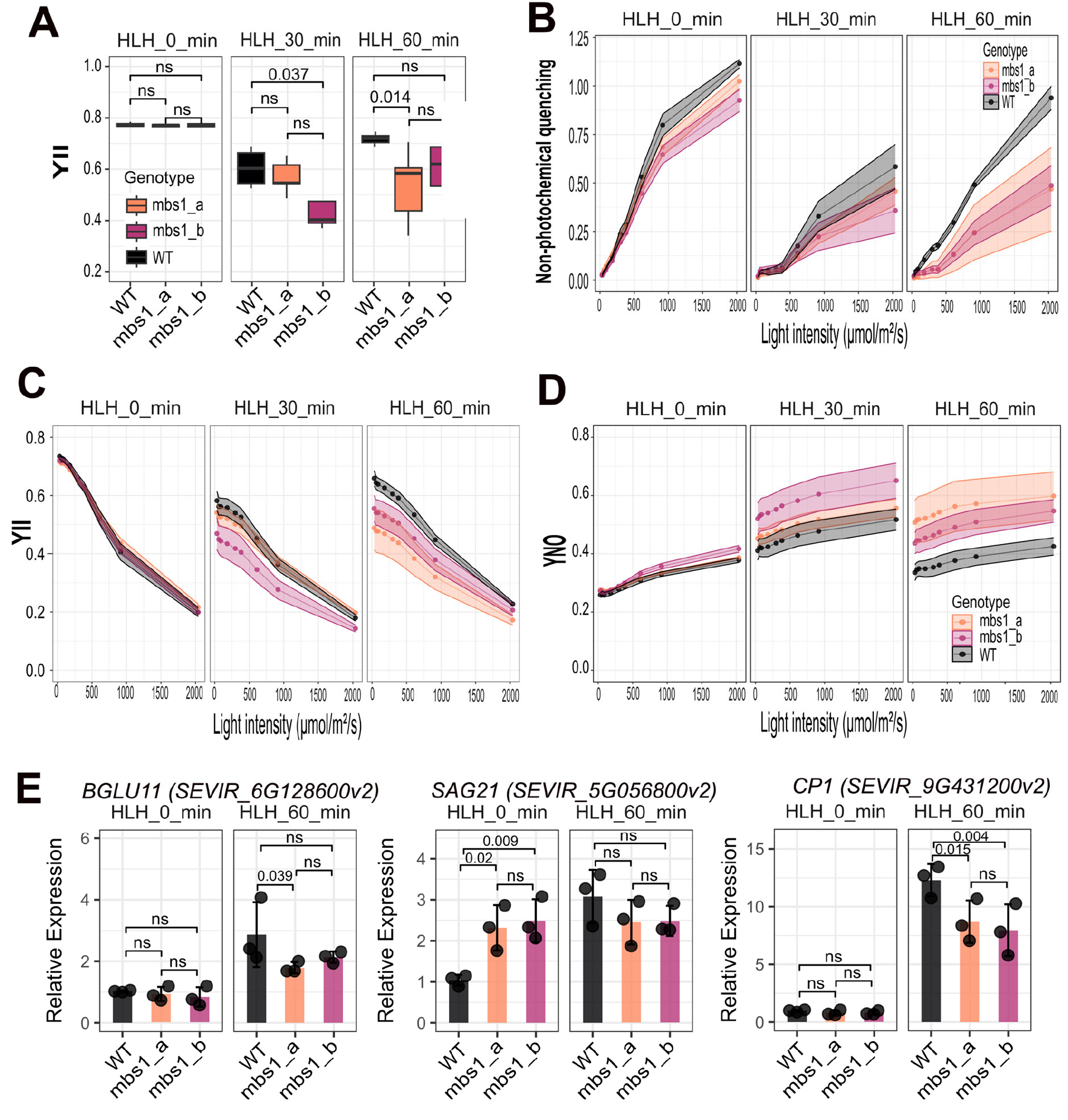
MBS1 is important for maintaining PSII photochemical efficiency, non-photochemical quenching, and stress responsive gene expression in mesophyll cells. **(A)** PSII maximum efficiency (*F_v_/F_m_*) measured by chlorophyll fluorescence in leaves of WT or 2 independent null *mbs1* mutant lines (mbs1_a and mbs1_b) after 0-, 30-, and 60-minutes HL+H treatment. Non-statistically significant differences between genotypes after a given duration of HL+H treatment are indicated by ns (Two-way ANOVA followed by Tukey’s post-hoc test, *p* < 0.05). **(B-D)** Light response curves of PSII photochemical and non-photochemical parameters measured via chlorophyll fluorescence. YII directly represents the effective quantum yield of PSII, NPQ the non-photochemical yield of PSII, and YNO the non-regulated non-photochemical yield of PSII, often indicative of photodamage as both photochemical energy conversion and NPQ are inefficient to convert or dissipate all light energy. **(E)** Expression of singlet oxygen-responsive marker genes in Mes sap extracted from WT or the 2 independent null *mbs1* mutant lines after 0- or 60- minutes of HL+H treatment as measured via real time quantitative PCR. Non-statistically significant differences between genotypes after a given duration of HL+H treatment are indicated by ns (Two-way ANOVA followed by Tukey’s post-hoc test, *p* < 0.05).

## DISCUSSION

There is an emerging hypothesis that chloroplast retrograde signalling performs specialised functions in accordance with chloroplast specialisation in different leaf cell types. However, most retrograde signalling research to date has been performed at the whole-leaf level. Only a few studies have considered individual cell types; but even so, these studies have focused on a single retrograde signalling pathway either in a single cell type, such as guard cells vs whole leaves^28^ or across cell types where the extent and nature of chloroplast specialisation remain unclear^29, 62^.

Here we systematically profiled the transcriptional networks associated with six stress-responsive chloroplast retrograde signalling pathways in C_4_ M and BS cells, two cell types where the specialisation of photosynthetic biochemistry in chloroplasts have been extensively detailed^38, 63^; and for two of these pathways, validated cell-specialisation of their signal metabolites and associated photosynthetic ROS. We show that three retrograde signals (DHA, PAP, DHAP) may function in a cell-type specific manner (**Fig. 6**). PSII ^1^O_2_-responsive DHA-MBS1 and PSI H_2_O_2_-responsive PAP signalling were restricted to M cells, consistent with the higher capacity for photosynthetically derived ^1^O_2_ and H_2_O_2_ production in M chloroplasts (**Fig 2,3**). In contrast, BS cells utilise rapid retrograde signalling via the Calvin cycle intermediate, DHAP, to activate AP2/ERF transcription factor-mediated transcriptional responses (**Fig 4**).

**Figure 6.**
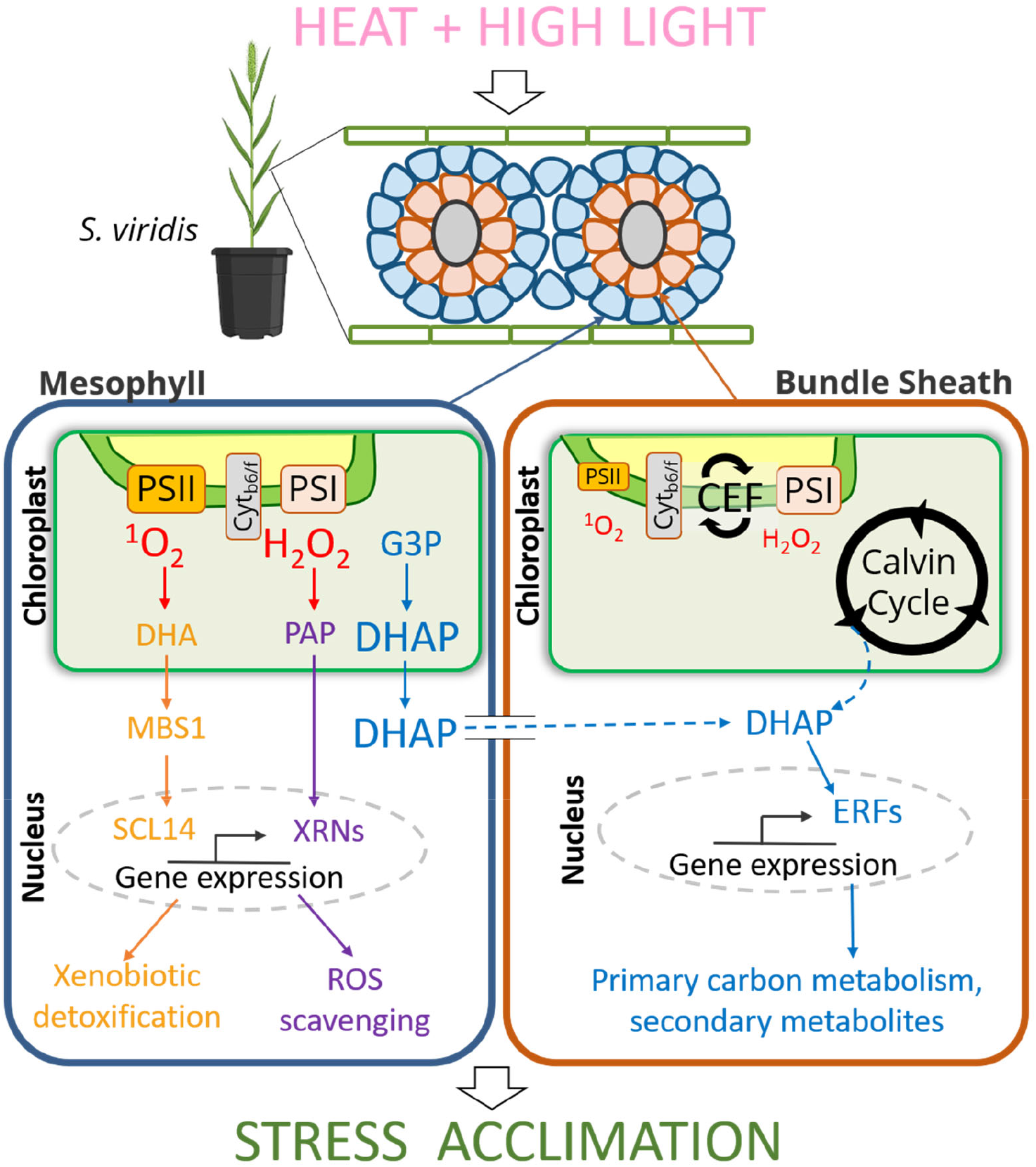
Specialisations of photosynthetic biochemistry potentiates cell-specific retrograde signalling in *S. viridis*. Schematic summarising the key cell-specific chloroplast-to-nucleus retrograde signalling pathways induced under high light and heat stress, as identified in this study via multi-omic analyses and functional characterisation of the selected ^1^O_2_-signalling mutant, *mbs1*. PSII-derived ^1^O_2_-responsive signalling is represented by the orange pathway in mesophyll cells. Mesophyll-specific PSI-derived H_2_O_2_-responsive retrograde signalling is represented by the purple pathway. BS-specialised chloroplast-to-nucleus retrograde signalling proposed to occur via DHAP is represented by the blue pathway.

### PSII enrichment in NADP-ME C_4_ mesophyll chloroplasts enable mesophyll-specific ^1^O_2_-dependent chloroplast signalling via DHA and *MBS1*

Plants performing the NADP-ME subtype of C_4_ photosynthesis show significant reduction in, or complete absence of, active PSII supercomplexes in BS chloroplasts^64^. We found that this is associated with significant loss of capacity for PSII-derived retrograde signalling in NADP-ME BS cells. *S. viridis* BS produced less ^1^O_2_ in their thylakoids, lacked stress-responsive accumulation of the ^1^O_2_ retrograde signal DHA, and failed to up-regulate the expression of ^1^O_2_-dependent marker transcripts implicated in xenobiotic detoxification (**Fig 2**). The ^1^O_2_-responsive DHA signalling was specialised to M cells, correlating with presence of functional PSII in this cell type (**Fig 2**). Therefore, cell type-specialisation of photosynthetic electron flow can be directly linked to a corresponding specialisation in retrograde signalling.

Although DHA and β-CC are both high light-induced and fulfil redundant functions in ^1^O_2_ signalling in *Arabidopsis*^60, 61^, only DHA, but not β-CC nor its downstream metabolite β-CCA, accumulated in response to HL+H treatment in *S. viridis* (**Fig 2D**). For β-CCA, this is likely because it is rapidly degraded under high light stress^65^. Oxidative cleavage of the double bond of the β-carotene polyene chain produces β-CC and β-ionone; the latter is oxidized by ^1^O_2_ to yield DHA^4^. β-ionone can also be cleaved by carotenoid cleavage dioxygenases to produce DHA^4^.The multiple routes for DHA production may explain its greater accumulation compared to β-CC in oxidatively stressed *S. viridis*, consistent with prior observations in high light-stressed *Arabidopsis*^66^ and suggests that DHA may be the more biologically relevant apocarotenoid signal in NADP-ME C_4_ plants.

*MBS1* has been extensively described as a signal transducer downstream of both DHA and β-CC^60, 67, 68^. In *Arabidopsis*, the MBS1 protein functions in stress acclimation by participating in liquid-liquid phase separation^67^. Null *mbs1* plants are insensitive to exogenous β-CC treatment and are photosensitive^60^. The stress-protective role of MBS1 is largely conserved in *S. viridis*, where WT SvMBS1 protein aggregated under HL+H treatment (**Fig S16**). The *mbs1* mutants had lower PSII activity after HL+H treatment, impaired induction of NPQ, and sustained photodamage, consistent with their hampered induction of M-specific gene expression (**Fig 5**). Interestingly, *MBS1* is expressed in both M and BS despite ^1^O_2_-derived DHA signalling being M-specific (**Fig S15, Fig 2**). This may reflect the deep evolutionary conservation of MBS1, which also functions in ^1^O_2_ signalling in algae^68^ and may offer stress-protective roles independent of chloroplast signals^67^, compared to the more recent evolution of NADP-ME C_4_ photosynthesis^32^.

It is possible that BS cells still retain ^1^O_2_-responsive signalling via the EX1/EX2 pathway. This pathway is spatially distinct from β-CC-and DHA-mediated signalling, being localised to the grana margins rather than the appressed thylakoids, which are absent from BS chloroplasts^6, 8, 69^. Dogra and Kim (2020) proposed that EX1 functions uniquely to sense ^1^O_2_ produced during PSII repair and biogenesis under more extreme oxidative stress conditions. Indeed, we found *EX1* to be more highly expressed in BS strands, which together with the BS-specific up-regulation of EX1 signalling markers such as *WRKY40* and *WRKY70* under HL+H, could reflect capacity for EX-dependent ^1^O_2_ signalling in BS (**Fig S10**).

### SAL-PAP chloroplast signalling is compartmentalised to mesophyll cells in *S. viridis*

SAL-PAP signalling has been proposed to drive differential cellular outcomes specific to different cell types^12, 28^. Foliar PAP accumulation in the Arabidopsis *sal1* mutant induces constitutive *APX2* expression, suppressing ROS accumulation in vascular bundles^12^. In contrast, exogenous PAP treatment induces ROS accumulation in both chloroplasts and nuclei of guard cells, leading to stomatal closure^28^. The BS cells of NADP-ME type C_4_ plants are enriched in PSI, the key origin of plastid H_2_O_2_ necessary for PAP accumulation, relative to M cells. Despite this, thylakoid membranes from BS cells produced less PSI-derived ROS than those of M cells (**Fig 3B**), likely due to the strong CEF in BS chloroplasts, which decreases electron leakage for H_2_O_2_ production at PSI^39, 70^. This may explain the lack of HL+H-responsive PAP accumulation in BS strands (**Fig 3C**), given that the PAP catabolic enzyme, SAL1 is inactivated by oxidation^11^. We identified two orthologous *SAL* genes in *S. viridis*, both of which were significantly enriched in BS strands relative to M sap **(Fig S9**). These factors may cumulatively enable stronger PAP degradation capacity in BS, thus preventing PAP accumulation in this cell type.

In M cells, PAP accumulated significantly under HL+H. PAP levels strongly correlated with the M-specific expression of established PAP marker transcripts such as *ELIP1*, *APX*, and *MDAR6* (**Fig 3A,E**), suggesting SAL-PAP signalling is M-specific in *S. viridis*. This was surprising because cell-specific SAL-PAP signalling was proposed to occur in BS^30, 71^ in conjunction with elevated *APX2* and *ZAT10* expression in Arabidopsis vascular bundles^72, 73^. In *Brassicaceae* species, such as *Arabidopsis*, glucosinolate biosynthesis is thought to be a major source of PAP^71^. As glucosinolate biosynthetic enzymes, including the PAP-producing SOTs, and SAL1 are highly expressed in *Arabidopsis* vascular bundles^74^, and there is significantly lower BS ROS accumulation in PAP-accumulating mutants^12^, SAL-PAP signalling is thought to be enriched in BS cells^30^. However, non-*Brassicaceae* plants lack glucosinolate biosynthesis, which is relatively enriched in specialised cells of *Arabidopsis* vascular bundles. Consequently, the balance between SOT-mediated PAP biosynthesis and SAL-mediated PAP degradation may be significantly different in non-*Brassicaceae*, enabling the emergence of M-specific PAP signalling. This may also explain the 10-20 fold PAP increase under HL (**Fig S2**), which is an order of magnitude higher that observed for HL treated Arabidopsis leaves^12^. Understanding the function of the three M-specific SOTs (**Fig S9**) and the two *SAL* orthologues during stress acclimation will contribute to testing this speculative hypothesis.

### The DHAP-AP2/ERF rapid retrograde signalling pathway is specific to BS cells

Rapid chloroplast retrograde signalling in *Arabidopsis* is dependent on export of the Calvin cycle intermediate DHAP from chloroplasts via the triose phosphate translocator (TPT), which then activates expression of various AP2/ERF transcription factors^19^. The rapid DHAP-AP2/ERF pathway is activated in the timescale of seconds to minutes and precedes ROS accumulation^19, 21^. These features are conserved in *S. viridis* as shown by the up-regulation of AP2/ERF-responsive transcripts within 30 min HL+H, and their down-regulation to near-unstressed levels by 60 min HL+H (**Fig 4E**). However, in C_3_ plants, M cells are the major sites of CO_2_ fixation via the Calvin cycle and thus the DHAP-AP2/ERF retrograde pathway would be expected to be most active in this cell type. In contrast, our data herein show that C_4_ plants perform BS-specific DHAP-AP2/ERF signalling (**Fig 4**). This provides a mechanism for BS cells to respond to abiotic stress in the absence of PSII ^1^O_2_ and PSI H_2_O_2_ signalling pathways in this cell type (**Fig 2, 3**); and may enable pre-acclimation of the BS to subsequent ROS bursts by the BS-enriched NADPH oxidases at the plasma membrane^75^.

In C_3_ photosynthesis, increased illumination causes an initial stimulation of photosynthetic light reactions and the Calvin cycle, thereby enhancing production of DHAP^76, 77^ and activation of AP2/ERFs^19^. Given that the Calvin cycle is localised exclusively to the BS in C_4_ plants33, BS-specific DHAP-AP2/ERF signalling could be expected. However, in C_4_ plants the M cells are thought to be the major source of the DHAP required in BS cells, with DHAP being shuttled between these two cell types via plasmodesmata^31^. Indeed, it is estimated that an approximately 10 mM gradient of DHAP between M and BS cells is necessary to support import of reduced carbon to BS chloroplasts^78, 79^. This raises two questions: 1) why is DHAP-AP2/ERF signalling inactive in M cells despite their substantially higher DHAP content; and 2) is the AP2/ERF pathway in BS cells triggered by DHAP from M cells, the BS Calvin cycle, or both? Resolving these questions will require identifying the as-yet-unknown DHAP receptor(s) and their cell-type expression, as well as understanding the cellular components regulating DHAP flux between M and BS cells during abiotic stress.

### Not all retrograde signalling pathways are cell type-specific

Transcripts for signalling components and marker genes of the EX1/EX2, GUN, and MEcPP retrograde signalling pathways were broadly expressed in both M and BS cells (**Fig S10-12**), suggesting that these pathways are not cell type-specific. This finding underscores the impact of investigating multiple retrograde signalling pathways in parallel as undertaken herein. A focus on individual pathways, as typically performed in retrograde signalling studies, would have led to significantly different conclusions on whether cell-specific retrograde signalling exists, depending on the pathway being studied.

Why are the EX1/EX2, GUN and MEcPP pathways expressed in both cell types? In the case of EX1/EX2, this may reflect the residual amounts of functional PSII in BS chloroplasts of NADP-ME *S. viridis* when grown at lower light intensities^64^, such as those employed in this study (250 µmol photon m^-2^ s^-1^). The GUN pathways are involved in chloroplast protein homeostasis and regulation of photosynthetic light reaction components, such as *LHCB*^23, 80^. Similarly, the MEcPP metabolite is derived from the MEP pathway, which feeds into chlorophyll biosynthesis^81^ and is expected to occur in both M and BS chloroplasts. MEcPP is involved in the regulation of several stress-responsive cellular processes including the ER unfolded protein response and hormonal homeostasis^17, 18^. The centrality of these processes to chloroplast function and photosynthesis may explain why the GUN pathway components are differentially expressed under HL+H treatment in both M and BS, albeit with some cell type-specific differences in how these transcripts are regulated (**Fig S12**) and with the caveat that GUN1 is also regulated at the post-translational level in *Arabidopsis*^27^.

The co-occurrence of GUN and MEcPP pathways, which both impact on protein homeostasis, in multiple cell types may explain why the core set of HL+H-responsive DEGs common across all cell types were enriched in GO terms relating to protein folding and general RNA metabolism (**Fig S4**). A broader question is how the diverse retrograde signalling pathways in common or unique to M and BS cells collectively influence cellular and whole-plant stress acclimation. For instance, MEcPP and PAP have been proposed to function in concert^82^. In *S. viridis* this MEcPP-PAP interaction would only occur in M cells, not BS cells, which may contribute to the M- and BS-specific subsets of MEcPP responsive genes (**Fig S11B**).

### Photosynthetic biochemistry as a driver for sensory plastid function?

It has been proposed that chloroplasts and plastids of certain cell types may function as ‘sensory plastids’ for sensing changes in environmental conditions and to initiate cell type-specific retrograde signalling^30^. This hypothesis was based in part on the assumption that BS chloroplasts and epidermal pavement cell plastids are not photosynthetic and therefore have likely gained new primary roles in signalling^30^. However, here we show that both M and BS chloroplasts perform cell type-specific retrograde signalling alongside their role in photosynthesis (**Fig 2-5**). We also did not identify evidence for any of the retrograde signalling pathways being specifically enriched or lost in epidermal peels (**Fig 2-4**). This may reflect sample preparation artefacts, as due to the nature of E peel sample preparation, cells from the abaxial epidermis are selectively enriched and we were unable to individually isolate pavement and guard cells. Alternatively, the C_3_-type photosynthetic pathway may be present in E tissue, as starch grains have been observed in guard cells and pavement cells of C_4_ plants, including NADP-ME maize^83, 84^. Collectively our data indicate that cell type-specialisation of chloroplast retrograde signalling pathways is facilitated by specific components of photosynthetic biochemistry (**Fig 2-4**), rather than photosynthetic capacity *per se*.

Our data indicates that presence of a photosynthetic component within a cell type on its own is not a predictor for signal initiation and retrograde signalling. Despite PSI being present in both M and BS, the EPR signal for PSI-derived ROS is substantially stronger in M thylakoids (**Fig 3B**), correlating with M-specific PAP accumulation and downstream PAP transcriptomic responses (**Fig 3C-F**). This demonstrates the potential for cell-specialisation of retrograde signalling to occur at multiple points from photosynthetic biochemistry and signal initiation to the signal’s accumulation, perception and response. Importantly, the cell-specialisation can also occur differently for different retrograde pathways, since the PSI ROS-responsive MEcPP appears functional in both M and BS (**Fig S11**).

The evolution of C_4_ photosynthesis involved extensive adjustments in chloroplast biochemistry, number, cellular positioning, and interactions with other organelles, such as peroxisomes and mitochondria^32, 34^. These alterations in chloroplast function may have necessitated the utilisation of specific retrograde signalling pathways tailored to each photosynthetic cell type. For instance, DHA and PAP signalling may be required in M cells due to the higher capacity of M thylakoids to produce ^1^O_2_ and H_2_O_2,_ respectively (**Fig 2-3**), compared to BS cells, where the higher CEF evolved to fulfil additional ATP requirements for C_4_ photosynthesis may already confer photoprotection^39, 70^. Whether the cell-specific chloroplast signalling observed herein in NADP-ME *S. viridis* is also evolutionarily conserved in other C_4_ subtypes where BS chloroplasts contain functional PSII, will require extensive investigation. Furthermore, an open question is how the cell type-specificity of retrograde signals are maintained in C_4_ plants where M and BS cells share extensive plasmodesmatal connections^40^. As plasmodesmata are regulated by chloroplast signals^85, 86, 87^, the interaction between plasmodesmata function and the chloroplast retrograde signals at both the M and BS interfaces will require investigation.

In conclusion, here we show that specialisation of photosynthetic biochemistry potentiates differential cell-specific retrograde signalling networks for distinct cell-specific stress acclimation (**Fig. 6**). This opens new avenues for understanding the co-evolution of cellular signalling networks and plant function^88, 89^, and identifies new considerations for the successful engineering of C_4_ photosynthesis into C_3_ plants for improved crop productivity^90, 91^.

## AUTHOR CONTRIBUTIONS

KXC, RTF and BJP designed the project; SY, RF, GB, KXC, NB and NC planned the experiments; SY, RF, SYP, NM and MTN performed experiments, SY, RF, NM and GB performed data analysis; SY, RF and KXC wrote the manuscript with input from all authors.

## ACKNOWLEDGEMENTS

We thank Dr. Maria Ermakova for sharing the hpt selection cassette and SpCas9 expression construct. We thank the Biomolecular Resource Facility (BRF) at the John Curtin School of Medical Research (JCSMR), Australian National University (ANU) for performing the Illumina cDNA library sequencing, and the Joint Mass Spectrometry Facility (JMSF), ANU for providing access to the gas chromatography-mass spectrometry instruments and research infrastructure used in this work. We acknowledge the use of the facilities, and scientific and technical assistance of the Australian Plant Phenomics Network (APPN), which is supported by the Australian Government’s National Collaborative Research Infrastructure Strategy (NCRIS).

## MATERIALS AND METHODS

### Plant Material and Growth Conditions

*S. viridis* plants (ME034V-1 ecotype) were germinated in individual 500 mL pots containing commercial seed raising mix and 3 g per L of Osmocote slow-release fertilizer. Plants were grown in controlled environmental chambers with a 16 h light/8 h dark photoperiod, 28 °C days, 24 °C nights, 60% humidity, and 185-200 μmol m^-2^ s^-1^ light intensity. For all experiments, plants were analysed 3-4 weeks after sowing, using the 3rd and 4th fully expanded leaves for harvesting or non-destructive measurements. HL+H, HH, or HL conditions were applied in a separate controlled growth chamber with temperature and light intensity raised to 39 °C and 1200 µmol m^-2^ s^-1^ light respectively. All experiments involving HL+H treatment were performed between the hours of 8:00-11:00 am to minimize the effect of circadian rhythm on physiology, metabolism and gene expression ^92, 93^.

WT *Nicotiana benthamiana* seeds were germinated in individual 135 mm (1.5 L) round plastic pots containing seed raising mix supplemented with Osmocote slow-release fertilizer, as described above. Plants were grown in a controlled temperature room under a 16 h light/8 h dark photoperiod, with temperature maintained at 21 °C, 55% relative humidity, and 325 μmol m^-2^ s^-1^ light intensity. Only the first, second and third true leaves from 4-5 week-old plants were used for agroinfiltration, while the rest were discarded.

### Generation of *S. viridis* lacking functional MBS1

*S. viridis* ME034V-1 plants with null *mbs1* alleles were generated via CRISPR/Cas9 gene-editing, as described by Ermakova *et al* ^94^. The genomic and coding sequences of *MBS1* (SEVIR_4G286500v2) were obtained from the *S. viridis* A10 v2.1 genome on Phytozome (https://phytozome-next.jgi.doe.gov). Two 20 nt guide RNAs (gRNAs) fitting the search criteria ‘A…19N…NGG’ within the *MBS1* single exon were selected using CRISPOR ^95^. Adenine (A) was required as the first base in the gRNAs to maximise expression by the *Oryza sativa* U3 snoRNA (OsU3) promoter^96^, while the NGG protospacer adjacent motif sequence was used for recognition by the *Streptococcus pyogenes* Cas9 (SpCas9) protein. The selected gRNAs (**Table S1**) were assembled according to Xie *et al* ^97^ into a synthetic polycistronic gene driven by the OsU3 promoter for processing via the endogenous tRNA-processing machinery. Further assembly into the plant binary expression vector, pICSL4723 was performed according to the Golden Gate (Type IIS) modular cloning guidelines established by Engler *et al* ^98^. The final plant expression vector carried the hygromycin phosphotransferase (hpt) gene driven by the O. sativa Actin 1 (OsAct1) promoter, and the SpCas9 gene driven by the Zea mays ubiquitin 1 (ZmUbi1) promoter.

The resulting construct was transformed into *S. viridis* ME034V-1 calli using the *Agrobacterium tumefaciens* strain AGL1 ^99^. T_0_ plants resistant to hygromycin were analysed for hpt copy number variation by qPCR (AttoDNA, Norwich, UK). Editing of *MBS1* was confirmed by Sanger sequencing (Macrogen, Korea) in segregated T_1_ plants without *hpt* and the progenies of plants with homozygous null alleles were used in all experiments. Two distinct null *mbs1* alleles were obtained and designated *mbs1 A* and *mbs1 B* (**Fig S16A**). Gene and protein sequences were visualized in Geneious Prime (Dotmatics, https://www.geneious.com/).

### Analysis of *mbs1* alleles via transient expression in *Nicotiana benthamiana*

CRISPR-edited *mbs1* sequences were amplified from crude DNA using the Phire Plant Direct kit (Thermo Fisher Scientific) and primers detailed in **Table S1**. Each of the two amplified *mbs1* alleles were adapted for modular Golden Gate cloning and cloned into the F2 acceptor vector (pICH47742)^100^, with expression driven by the ZmUBI1 promoter and terminated by the *A. tumefaciens* nos terminator sequence. An mCitrine reporter was also tagged onto the C-terminal region of each MBS1 isoform for visualisation. Assembled plasmids were transformed, amplified and maintained in the *Escherichia coli* Top10 strain prior to *Agrobacterium*-mediated infiltration for transient expression *in planta*.

C-terminally-tagged *mbs1* mutant isoforms were transiently expressed in *N. benthamiana*, following established protocols^101^. Briefly, *Agrobacterium tumefaciens* strain GV3101 (pMP90) transformed with the constructs of interest were cultured in LB media with 25 µg/mL rifampicin, 50 µg/mL gentamycin and 50 µg/mL carbenicillin at 28 °C and 220 rpm for 24 h. *A. tumefaciens* strain GV3101 (pMP90) containing a plasmid encoding the Tomato Bushy Stunt Virus P19 viral suppressor protein was also mixed with cultures containing the plasmid(s) of interest at optical densities (OD 600 nm) of 0.3 and 0.8, respectively for co-infiltration. A P19-only control was prepared to an OD_600_ of 0.8 as a negative control. Following overnight growth, bacterial cultures were pelleted at 2,150 g for 8 min and resuspended in infiltration solution (10 mM MES pH 5.7, 10 mM MgCl2, 150 μM acetosyringone). Resuspended cells were incubated at RT for 2 h prior to infiltration into the abaxial surface of 4-5 week-old *N. benthamiana* leaves using a needle-less syringe.

Transient protein expression and HL+H treatment was assessed 3 days post-infiltration (dpi) via widefield fluorescence imaging on the Leica DM5500 microscope. Briefly, *N. benthamiana* leaf disks were harvested from infiltrated spots, vacuum infiltrated with water for 1 min and mounted on microscopy slides with their abaxial surfaces facing up. To avoid crushing the leaf disks, cover slips were elevated with a small drop of petroleum jelly at each corner. Mounted slides were placed in the HL+H stress chamber detailed above for 0, 15 and 30 min prior to widefield fluorescence microscopy, as described in SI Appendix.

### Mechanical cell separation and enrichment

A detailed description of mechanical cell separation techniques can be found in SI Appendix. Briefly, M sap was isolated using leaf rolling as described in Covshoff *et al* ^37^, BS strands were isolated via blending and differential filtration as described in John *et al* ^46^, and E peels prepared according to Weyers and Travis^102^.

### RNA extraction

Total RNA was isolated from cell type-enriched fractions using the RNeasy Plant Mini Kit (Qiagen), following a modified protocol. Briefly, snap frozen BS strands and E peels were disrupted with steel balls using a TissueLyser II (Qiagen) at 25 Hz for 1 min. To each powered sample, 450 μL Buffer RLT supplemented with 40 mM DTT was added. M sap samples were extracted into 450 µL of Buffer RLT instead, as outlined in the SI Appendix. All lysates were homogenized through the QIAShredder spin columns before mixing with half volume of 100% ethanol and transferred to RNeasy Mini spin columns. Crude RNA collected on the columns was washed with 500 μL of Buffer RW1, followed by on-column DNase treatment with 3 U of amplification grade DNase I (Invitrogen) for 15 min. Columns were then washed once with 700 μL of Buffer RW1 and twice with 500 μL of Buffer RPE (containing 80% ethanol) to remove impurities before spin-drying. Purified RNA was eluted with 25-30 μL of nuclease-free water. For samples with expected low RNA yields, such as from E peels, elution was repeated using the initial eluate to increase the concentration of recovered RNA.

Total RNA was quantified spectrophotometrically at 260 nm using a NanoDrop1000 Spectrophotometer (Thermo Fisher Scientific). RNA quality was assessed via capillary electrophoresis using a LabChip GX Touch Nucleic Acid Analyser (Perkin Elmer) with a high throughput RNA Pico Sensitivity Assay reagent kit (Revvity).

### Reverse Transcription -quantitative PCR (RT-qPCR)

cDNA synthesis was performed using 1000 ng of total RNA according to the SuperScript IV First-Strand Synthesis System (ThermoFisher Scientific). 10 ng cDNA was subjected to qPCR analysis in the CFX Opus 384 Real-Time PCR System (Bio-Rad), using the SensiFAST SYBR Lo-ROX Kit (Bioline) according to manufacturer’s instructions. Calculation of relative transcript levels were performed using LinRegPCR and data normalised to the housekeeping gene *CUL* and to WT control tissue^103, 104^

### Sequencing library preparation and analysis

mRNA sequencing libraries were prepared using the TruSeq mRNA Stranded Kit (Illumina) with minor modifications, including scaling reactions to one-third, as detailed in Crisp *et al*^13^. The amount of starting total RNA used was standardised across all samples for consistency and normalisation. Specifically, 1.8 μg of total RNA was used to generate the time-resolved, cell type-enriched bulk RNA-seq dataset in this study. First-strand synthesis was performed using SuperScript III, followed by library clean up and size selection using Sera-Mag Solid-Phase Reversible Immobilisation (SPRI) beads. Fragments that were approximately 380 bp in length (250 to 700 bp) were typically selected for. Adapter ligation was performed using the IDT for Illumina TruSeq DNA Unique Dual (UD) Indexes kit, which enables dual indexing with i5 and i7 regions. Libraries were PCR-amplified for 10 cycles, based on the starting quantities determined using the dsDNA High Sensitivity Assay kit (Invitrogen) on the Qubit 2.0 Fluorometer (Invitrogen). Individual and pooled library sizes were analysed using a high throughput DNA High Sensitivity reagent kit (Revvity) on the LabChip GX Touch NucleicAcid Analyser (Perkin Elmer). Libraries were pooled in equimolar ratios to achieve a final concentration of 2.5 nM for sequencing.

The pooled cDNA library was sequenced on an Illumina NovaSeq 6000 platform at the Biomolecular Resource Facility (BRF), Australian National University. Libraries were sequenced using an S2 flow cell with 150 bp paired-end sequencing and loaded at 300 μL of 2.5 nM pooled cDNA, yielding 1000-1250 Gb (3.3-4.1 billion reads) per flow cell across 96 multiplexed samples (approximately 10.4-13.0 Gb or 34-43 million reads per sample). All samples were run on a single lane to avoid batch effects.

Quality control of raw sequencing reads was performed using FastQC. Adapters and polyA/T tails were removed using CutAdapt^105^ within TrimGalore, with a quality threshold (Phred score, Q) of 20. A HISAT2^106^ index containing exons and alternative splice sites was generated from the *S. viridis* A10 v2.0 reference genome^107^ obtained from Phytozome (v13, https://phytozome-next.jgi.doe.gov/). Trimmed reads were aligned to this index using HISAT2, with parameters specified for paired-end reads and strand-specific information to identify reads mapping to positive and negative strands of the genome. The resulting sequence alignment files were processed using SAMtools^108^ for sorting, merging, indexing, compressing, and converting from Sequence Alignment/Map (SAM) to Binary Alignment/Map (BAM) format. Fragment counts per gene were calculated using featureCounts^109^, with the -p and --countReadPairs flags specified to count and summarise uniquely mapped read pairs comprising a fragment that map to exons within each gene locus.

### RNA-seq Data Processing, Differential Expression, and Network Analysis

Paired-end RNA-seq reads uniquely mapped to the *S. viridis* A10 v2.0 reference genome were processed prior to differential gene expression analysis. Specifically, filtering to remove lowly expressed genes with a row sum of < 25 in 5 or more samples was performed to reduce noise, in addition to removing genes encoded within the chloroplast and mitochondrial genomes. All read counts were normalized using DESeq2, and DEGs identified using a stringent threshold of adjusted *p* value < 0.05 and a log2 fold-change (LFC) either > 1 or < -1 to define differential expression^47^. *K*-means clustering was performed according to Hartigan and Wong ^110^ GO analysis using the g:Profiler2 package, and heatmaps produced using the ComplexHeatmap package. Co-expression network analysis was performed according to Li *et al* ^48^, and GRNs constructed using the Genie3 package and visualized in either Cytoscape or the R package igraph. For *Cis-*regulatory element analysis, promoter sequences (1.5-kb upstream of the predicted start codon) were obtained from Ensembl, and subject to *cis*-regulatory enrichment analysis using Regulatory Sequence Analysis Tools (RSAT) web server^111^.

### Thylakoid Isolation and EPR spectroscopy

Thylakoids from M and BS cells were isolated as described in Ermakova *et al* ^112^. For EPR spin-trapping experiments, thylakoids were suspended in activity buffer (20 mM MES-KOH, 10 mM EDTA, 400 mM sucrose, 15 mM NaCl, pH 6.5) to chlorophyll concentration of 10 µg per mL and incubated with either DMPO or TEMPD (Sigma-Aldrich, St. Louis, USA) at a final concentration of 50 or 100 mM respectively. Samples were loaded into 100 µL quartz flat cells and the baseline EPR signal acquired in the dark, before illuminating with 1200 μmol m^-2^ s^-1^ light for 2 or 10 minutes for DMPO and TEMPD experiments respectively. EPR measurements were acquired at room temperature with a model no. ELEXYS E500 Spectrometer (Bruker, Billerica, MA) equipped with an ER 4103TM resonator operating at X-band frequencies (9.5 GHz) and a 100 kHz modulation frequency. The following parameters were used: microwave frequency, 9.73 GHz; modulation frequency, 100 kHz; modulation amplitude, 1 G; microwave power, 6.3 mW ; time constant, 40.96 ms; and number of scans, 5.

### β-cyclocitral and dihydroactinidiolide quantification via SPME-GCMS

Solid-phase microextraction followed by gas chromatography and mass spectrometry (SPME-GCMS) was performed as described by Rivers *et al* ^51^ to quantify the accumulation of volatile organic compounds. Detailed information can be found in SI appendix.

### Adenosine Extraction and Quantification via HPLC

Total plant adenosines including PAP were extracted, derivatized with chloroacetaldehyde (CAA) and resolved by reverse phase high-performance liquid chromatography for fluorometric quantification as described in Bowerman *et al* ^113^ which is a method modified from Estavillo *et al* ^12^, and Bürstenbinder *et al* ^114^ . Detailed information can be found in SI appendix.

### Chlorophyll quantification

Total chlorophyll content of leaves, BS strands, and M sap were determined using aliquots set aside during metabolite extraction. Ground plant samples in either 2:1 chloroform methanol or 20% NaCl were diluted 1:10 in methanol prior to loading into 96-well plates. Absorbance of the samples at 652, 665.2, and 750 nm made measured an Infinite M-1000 pro Microplate Reader (Tecan) and calculated according to Porra *et al*^115^.

### Dual Pulse-Amplitude-Modulation (PAM) Fluorometry

Simultaneous measurements of chlorophyll fluorescence and the redox state of P700 (the PSI reaction centre) were used to determine quantum yields of PSII and PSI, using a Walz DUAL-PAM/F under red actinic light of 635 nm, and saturating pulses of red light (300 ms, 12,000 μmol m^-2^ s^-1^). Detailed information can be found in SI appendix.

### Data visualisation and statistical analysis

Data visualisation, processing, and statistical analyses were performed in R (version 4.3.X) with R Studio. Statistical tests for differential gene expression analyses are described in the relevant sections above.

