## Supplementary Methods and Figures for "Specialisation of photosynthetic biochemistry in C_4_ chloroplasts potentiates distinct cell-specific functionality of different chloroplast-to-nucleus retrograde signalling pathways"

This PDF file includes:

Extended Materials and Methods

Supporting References

Figures S1 to S16

Table S1

#### EXTENDED MATERIALS AND METHODS

##### Mechanical Cell Separation and Enrichment

After 0, 30 or 60 minutes of stress treatment, the mid-sections of 3rd and 4th fully expanded leaves of plants were harvested. E peels were prepared by making an incision on the adaxial surface of the leaf, taking care not to cut through the abaxial epidermis. Using the remaining leaf tip as a tab, the abaxial layer of E cells was gently peeled using forceps while securing the adaxial layers. The abaxial epidermal layers were immediately snap frozen in liquid $N_2$.

M sap was extracted via leaf rolling. First, the mid-ribs were removed for the mid-sections of harvested leaves and discarded. Leaf strips were then transferred to a pre-chilled square petri dish on ice and rolled following the direction of the veins using a Western blot roller to squeeze out M sap. In the case of preparing samples for PAP extraction, this sap was rolled into 300 µL of chilled citric acid-sodium phosphate (CP) buffer (61.45 mM citric acid, 77.1 mM $Na_2HPO_4$, pH 4.0). For β-CC and DHA quantification, M sap was rolled into 2 mL of chilled 20% NaCl, and for RNA extraction into 450 µL of Buffer RLT from the RNeasy Plant Mini kit supplemented with 40 mM DTT before freezing in liquid $N_2$.

To isolate BS strands, the mid-sections of the 3$^{rd}$ and 4$^{th}$ fully expanded leaves of 5-6 plants per sample were transferred to 15 mL of phosphate-sorbitol extraction buffer (50 mM sodium phosphate buffer pH 7.5, 330 mM sorbitol, 2 mM EDTA) and sliced into approximately 3 mm wide strips, before transferring with all liquid into a pre-chilled Omnimixer chamber and blended (3 bursts of 10 seconds each at speed 7) in an Omnimixer homogenizer (Omni International). This was then passed through a 500 µm mesh filter, and any solid debris blended again with an additional 15 mL of extraction buffer. Filtering through the 500 µm filter was repeated before washing the Omnimixer chamber with an additional 15 mL of cold extraction buffer and passing this through the 500 µm mesh filter. All filtrates were then pooled and passed through a 70 µm mesh filter to collect crude BS strands, which were then washed with 10 mL of cold extraction buffer. BS strands were collected by inverting the filter and washing with 2-4 mL of cold extraction buffer, eluting into an Eppendorf tube. These were then centrifuged at 21,000 *g* for 5 minutes at 4 °C, before removing supernatant and flash freezing in liquid $N_2$.

##### Chlorophyll fluorescence and P700 redox spectroscopy

The redox state of P700 (used for deriving PSI parameters) was monitored by detecting absorbance changes at 830 nm in the $P700^+$ cation. Plants were treated with 0, 30, or 60 minutes of HL+H prior to dark adaptation for 10 minutes, after which a saturating pulse was applied to determine the maximum ($F_M$) and minimum ($F_0$) PSII fluorescence in the dark. These were then used to calculate $F_V/F_M$, or the maximum quantum yield of PSII as $F_V/F_M = (F_M-F_0)/ F_M$ (Klughammer and Schreiber, 2008a)). After this, the maximum signal of $P700^+$ (PM) was determined by illuminating the leaf for 8-seconds with far-red light of 720 nm before applying a second saturating pulse. The minimal $P700^+$ signal ($P_0$) was recorded directly before this saturating pulse. Following the determination of these dark parameters, light response curves were performed by progressively increasing the intensity of actinic light from 55 to 1650 µmol $m^{-2}$ $s^{-1}$, illuminating the leaf for 100 s at the new light intensity before applying a saturating pulse. Upon each saturating pulse, the maximum PSII fluorescence under light ($F_M$\`), and the fluorescence level directly before the saturating pulse (F) were recorded. Using these values, the partitioning of the excitation energy absorbed at PSII between the photochemical and non-photochemical reactions could be tracked. This allowed for calculations of PSII photochemical yield ($YII = (F_M`-F)/F_M`$), the yield of regulated NPQ ($YNPQ = (F/ F_M`)-(F/F_M)$), and the yield of non-regulated NPQ ($YNO = F/F_M$), according to

Kramer et al. (2004) and Bilger and Bjorkman (1990). PSI parameters were derived using the steady-state $P700^+$ signal (P) and the maximum signal under light ($P_M$`) recorded just before the saturating pulse. This included the photochemical yield of PSI (YI = ($P_M$`-P)/($P_M$-$P_0$)), as described by Klughammer and Schreiber (2008b).

**Adenosine Extraction and Quantification via HPLC**

Unless specified otherwise, all steps were performed on ice or at low temperature to preserve metabolite samples. Snap frozen plant samples (E peels, BS strands, M sap) were ground at 25 hz for 1 minute using a TissueLyser II (Qiagen) with two 3 mm metal ball bearings. Samples were then centrifuged (1 min, 13,000 *g*, 4 °C) and then resuspended in 600 µL of chlorophorm:methanol (2:1 v/v). Samples were homogenised by Tissue Lyser again (25 hz, 1 min) and then 50 µL of the extract set aside for chlorophyll analysis. 300 µL of CP buffer was then added to each sample, before homogenising by Tissuer Lyser a final time (25 hz, 1 min), then centrifuging to separate solvent layers (5 min, 13,000 *g*, 4 °C). 250 µL of the hydrophilic upper phase was then transferred to a new Eppendorf tube. suitable for freeze drying. Samples were then snap-frozen in liquid N2 before lyophilisation under vacuum for 12-14 hours at -80 °C. Once lyophilisation was complete, samples were resuspended in 250 µL of dilute CP buffer (2:3 v/v, CP buffer:dH2O), before derivatizing all adenosines with 21.6 µL of 45% CAA (Sigma). CAA selectively reacts with adenosine rings to form "etheno compounds" that fluoresce at 410 nm upon excitation at 280 nm (Haink and Deussen, 2003). Samples were then centrifuged (5 min, 21,600 *g*), before heating at 80 °C for 10 minutes. Once heated, samples were cooled to room temperature on ice and centrifuged (45 min, 21,600 g), and the supernatant subjected to analysis via HPLC using a Hewlett-Packard/Agilent 1260 Infinity Series HPLC (Agilent Technologies) outfitted with a fluorescent detector. The gradient for separation of PAP was previously optimised and used the following HPLC run sequence: equilibration of Phenomenex Kinetex C18-XB column for 0.1 min with 95 % of buffer A [5.7 mM [CH3(CH2)3]4NHSO4, 30.5 mM KH2PO4, pH 5.8] and 5% buffer B [67 % acetonitrile, 33 % buffer A], linear gradient for 13.75 min up to 50 % of buffer B, re-equilibration for 1.66 min with 5 % buffer B. Adenosines were detected by fluorescence (excitation 280 nm and emission 410 nm). Alongside plant samples, commercial standards containing a mix of derivatized adenosine mono- (AMP, Sigma 4578-31-8), di- (ADP, Sigma 01905), and tri-phosphate (ATP, Sigma 213-579-1), PAP (Sigma, A5763), and 3'-phosphoadenosine-5'-phosphosulfate (PAPS, Sigma A1651) were run to aid identification of PAP in complex samples, as well as derivatized PAP standards of known concentrations (0, 0.04, 0.4, 1, 2, and 4 pmol $\mu L^{-1}$) to construct calibration curves for PAP quantification in each sample. Chromatograms were then processed with the Agilent LC1260 ChemStation software and integrated to quantify the area under the curve corresponding to the PAP peak in each sample. The calibration curve of PAP standards was then used to calculate the PAP content of each sample, before normalizing to chlorophyll content.

**Volatile apocarotenoid quantification via SPME-GCMS**

Frozen samples leaf and BS strand samples were ground to a fine powder by Tissue Lyser (25 hz, 1 min) and resuspended in 2 mL 20% (w/v) NaCl supplemented with 10 ppb of the internal standard 13,13,13-D3-β-ionone (Buchem V). 100 µL were set aside for chlorophyll analysis and the remainder of samples transferred to 20 mL amber vials. Samples of M sap were extracted directly in 2 mL 20% NaCl and transferred directly to amber vials. Samples were separated using a single quadrupole GC/MSD machine (Agilent technologies). The instrument consisted of a split/splitless injector and a 5975C inert XL MSD mass selective detector (Triple-Axis Detector), retrofitted with a MPS 2 Gerstel Multipurpose sampler with normal liquid injection and HS-SPME capability (Gerstel GmbH & Co. KG, Germany). All samples and standards were then heated (5 min, 70°C), before adsorbing volatile apocarotenoids onto a bipolar 50/30μm PDMS/DVB/CAR sorption SPME fibre for 40 min. Within the GC inlet, the fibre was heated again for 40 min to desorb volatile compounds, before using splitless injection with a 1.0 mm straight no-wool liner with the inlet temperature set to 250 °C. Volatile compounds were then separated on a non-polar Agilent J&W VF-5 ms column (30 m × 0.25 mm × 0.25 μm), equipped with a 10 m EZ-Guard column. The oven temperature programme was as follows: 40 °C for 2 min, ramping to 150 °C at 5 °C/min and held for 2 min, then to 320 °C at a ramp rate of 15 °C/min and held for 1 min. Helium (ultra-high purity, BOC Australia) was flowed at 1 mL/min, aux transfer lines heated to 320 °C, the ion source to 250 °C, and quadrupole to 150 °C. Electron impact ionisation energy (EI) was set to 70 eV, using full MS scan acquisition from *m/z* 40-500 with a 4 min solvent delay, giving a total GC/MS run time of 38 min.

For all data analyses the Agilent MassHunter software (B.50) was used. To quantify β-CC and DHA, their quantifier ion peaks (137 and 111 *m/z* respectively) in addition to that of the internal standard (180 *m/z*) were extracted from each chromatogram and the peak area quantified. B-CC and DHA content of plant samples was determined by comparing peak areas to those of pure β-CC standards of known quantities, adjusting for any differences in the abundance of the internal standard, prior to normalisation to chlorophyll.

**ROS staining and live cell imaging**

ROS were visualized in live tissue of *S. viridis* using the probe 2',7'-dichlorodihydrofluorescein diacetate ($H_2$DCFDA, Sigma D6883), which reacts with ROS to produce the fluorescent DCF. The mid-section of the fourth fully expanded leaves of plants were harvested and submerged into 10 mL of $H_2$DCFDA diluted to 500 μM in 50 mM sodium phosphate buffer (Na2HPO4.7H2O, Na2HPO4.H2O, pH 7.4) before vacuum infiltrating for 10 minutes or until vigorous bubbling subsided. Thin cross-sections of infiltrated leaves were then prepared by hand and washed in distilled water for two minutes in the dark before imaging on a Leica DM5500 fluorescence microscope (Leica microsystems) outfitted with a 480/40 bandpass excitation filter, 505 dichromatic mirror and 527/30 bandpass suppression filter. To confirm $H_2$DCFDA penetration into leaf tissue after infiltration with $H_2$DCFDA, thin sections of *S. viridis* leaves were prepared and immersed in 1 mL of 1 mM $H_2O_2$. After incubating in the dark for 5 minutes, these sections were washed in distilled water for two minutes prior to imaging.

All quantifications of fluorescence signal were made using the Leica Application Suite X. Total pixel intensity of regions of interest on the $H_2$DCFDA fluorescence channel were divided by the total region area. For each fluorescent micrograph, the background signal was also quantified using a fixed region of interest in the corner of the image, and this subtracted from any cell fluorescence quantifications.

**Widefield fluorescence microscopy of transiently expressed *mbs1* alleles exposed to HL+H treatment**

Widefield fluorescence microscopy was performed on transiently infiltrated *N. benthamiana* leaves expressing the mCitrine-tagged *mbs1* alleles. Images were captured on a Leica DM5500 fluorescence microscope (Leica Microsystems) fitted with a DFC7000T camera. All samples were imaged under a 40× water immersion objective with a sensor resolution of 1920x1440 pixels using the Leica Application Suite X (LAS X) software package. mCitrine signal was detected in *N. benthamiana* using a 500/20 bandpass excitation filter, 515 dichromatic mirror and 535/30 bandpass suppression filter. All image processing post-capture was performed in LAS X Office (version 1.4.5).

#### RNA-seq Data Processing, Differential Expression, and Network Analysis

*Co-expression and regulatory network construction*

The co-expression network was constructed using scaled and normalised variance stabilized expression data for all genes differentially expressed in response to HL+H in any cell type. An edge selection cutoff was defined using the Pearson correlation coefficient (adjusted for multiple comparisons using the Benjamini-Hochberg procedure) of > 0.85 to filter for highly positively correlated gene pairs. The co-expression network was then created using the graph_frome_data_frame() function of the igraph package. Co-expression modules were then detected using the Leiden algorithm, using an optimized resolution parameter of 3. The WGCNA package in R was then used to integrate information related to metabolite accumulation and perform module- and gene-level correlation analyses.

To construct the GRN using GENIE3, variance stabilised count data for all genes within the co-expression network was provided to the GENIE() function as the expression matrix, alongside the full list of *S. viridis* transcription factors and kinases obtained from the Plant Transcription Factor Database as potential regulators (Jin et al. 2017). This network was then filtered using via graph-based clustering implemented in the grn_filter() function, retaining approximately 2% of the original network edges. Both coexpression and regulatory networks were visualised using either Cytoscape (v 3.10.4) or the iGraph package in R.

*Enrichment analysis*

Gene ontology enrichment analysis was performed using the g:Profiler2 package in R. *S. viridis* DEGs were first mapped to their *Setaria italica* orthologues using the *gorth()* function, due to functional and gene ontology annotations being more robust and developed in this closely related species. GO term functional enrichment analyses were then performed using the *gost()* function, specifying a custom background containing all genes detected in the RNA sequencing experiment and using a Fisher's exact test with a false discovery rate cutoff of < 0.05.

### SUPPORTING FIGURES

#### Supplementary Figure 1

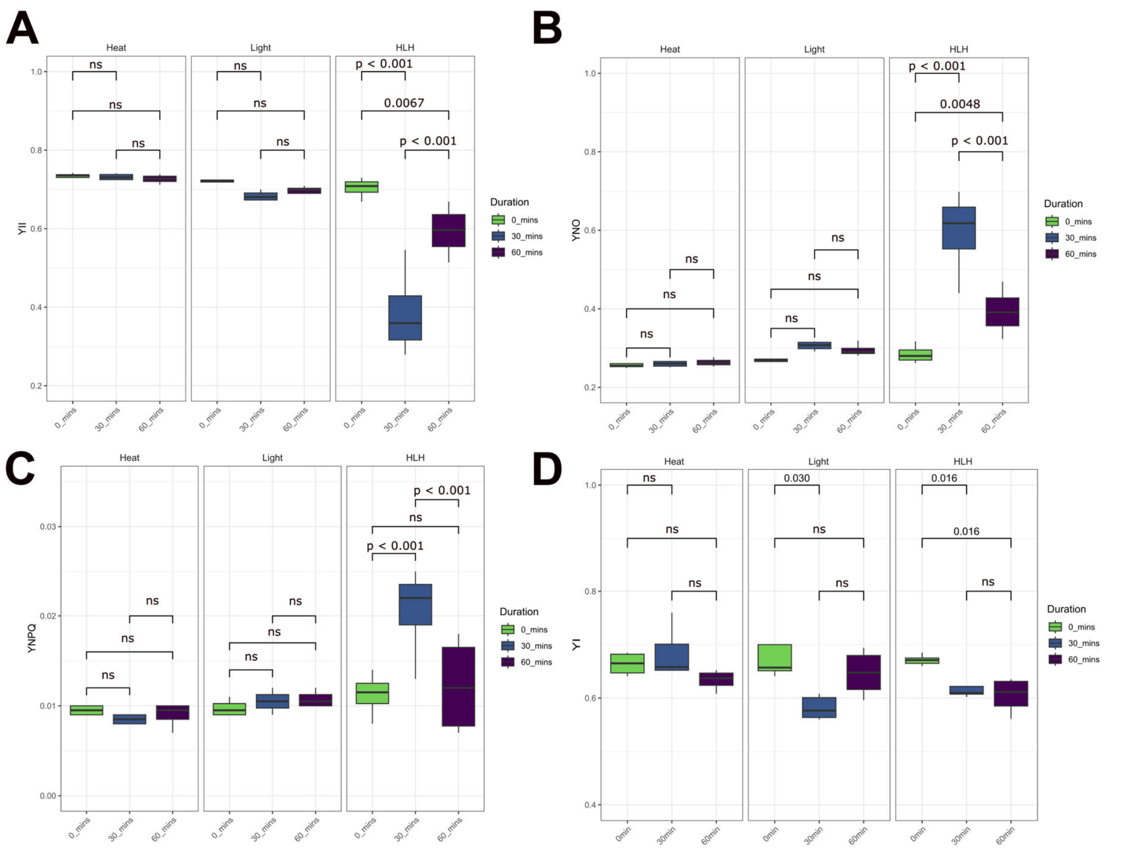


**Supplementary Figure 1. Combined high light and heat treatment significantly perturbs photosynthetic electron transport in *S. viridis* leaves.**
**(A-C)** Show photosystem II photochemical and non-photochemical yields simultaneously measured via chlorophyll fluorescence. **(A)** YII directly represents the effective quantum yield of PSII.
**(B)** YNO represents the non-regulated non-photochemical yield of PSII, often indicative of photodamage as both photochemical energy conversion and NPQ are inefficient to convert or dissipate all light energy.
(**C)** YNPQ represents the non-photochemical yield of PSII due to NPQ, or the energy dissipated from the PSII reaction centre as heat. **(D)** Quantum yield of photosystem I (YI) in *S. viridis* leaves after 0, 30, and 60 minutes of either heat (Heat), high light (Light), or combined high light plus heat treatment (HLH) as measured via $P700^+$ spectroscopy. Non-statistically significant differences between durations of stress treatment are indicated by ns (Two-way ANOVA followed by Tukey's post-hoc test, $p < 0.05$).

#### Supplementary Figure 2

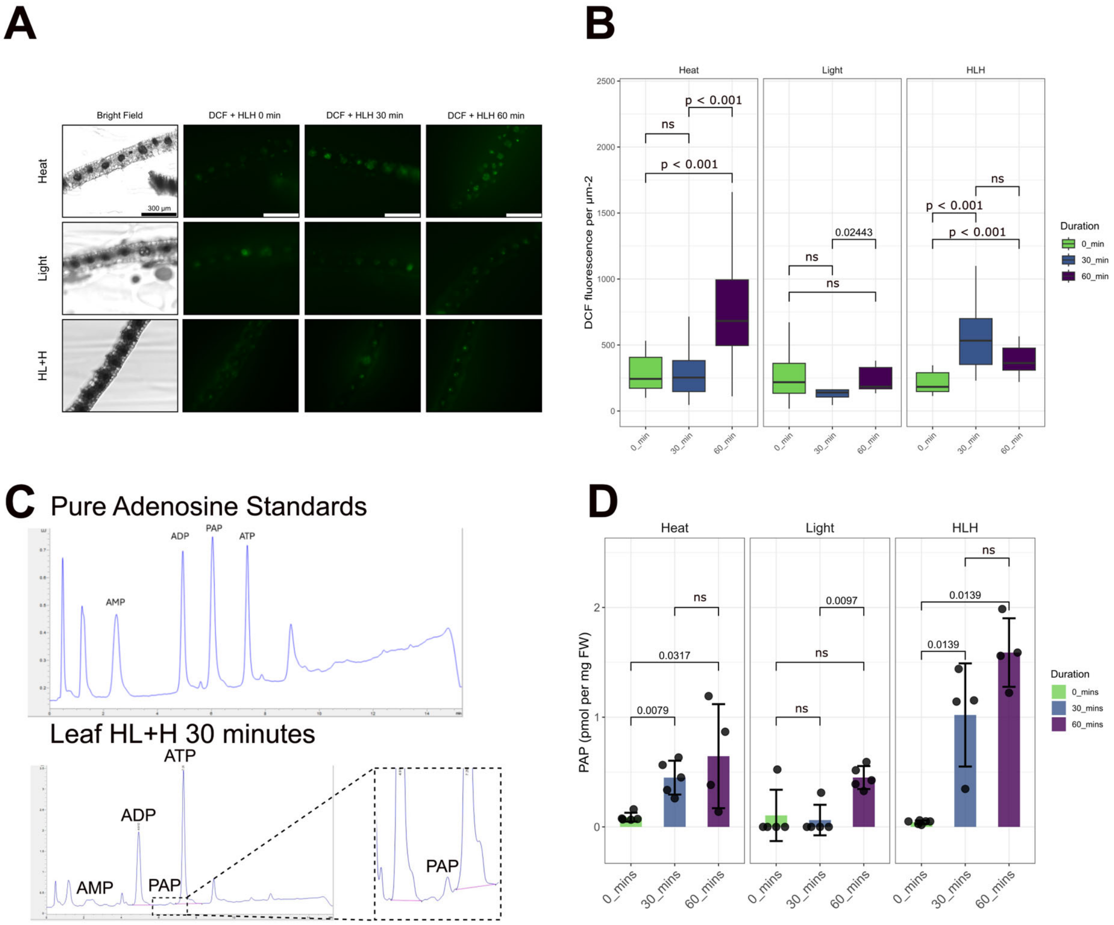


**Supplementary Figure 2. Combined high light and heat treatment induces significant ROS and chloroplast signal accumulation in *S. viridis* leaves.**
**(A)** Representative fluorescence micrographs showing ROS fluorescence via $H_2$DCFDA staining false-coloured green in cross-sections of *S. viridis* leaves after heat, light, or combined high light and heat treatment. White scale bars indicate 300 µm. **(B)** Quantifications of DCF fluorescence signal in *S. viridis* leaves after different stress treatments, expressed as pixel intensity per square µm. **(C)** Representative HPLC chromatogram of pure adenosine standards and a sample of mesophyll sap after 30 minutes of high light and heat treatment. **(D)** Absolute PAP levels (expressed as picomoles per mg of fresh weight) in *S. viridis* leaves as quantified by HPLC. For all experiments, non-statistically significant differences between durations of stress treatment are indicated by ns (Two-way ANOVA followed by Tukey's post-hoc test, $p < 0.05$).

#### Supplementary Figure 3

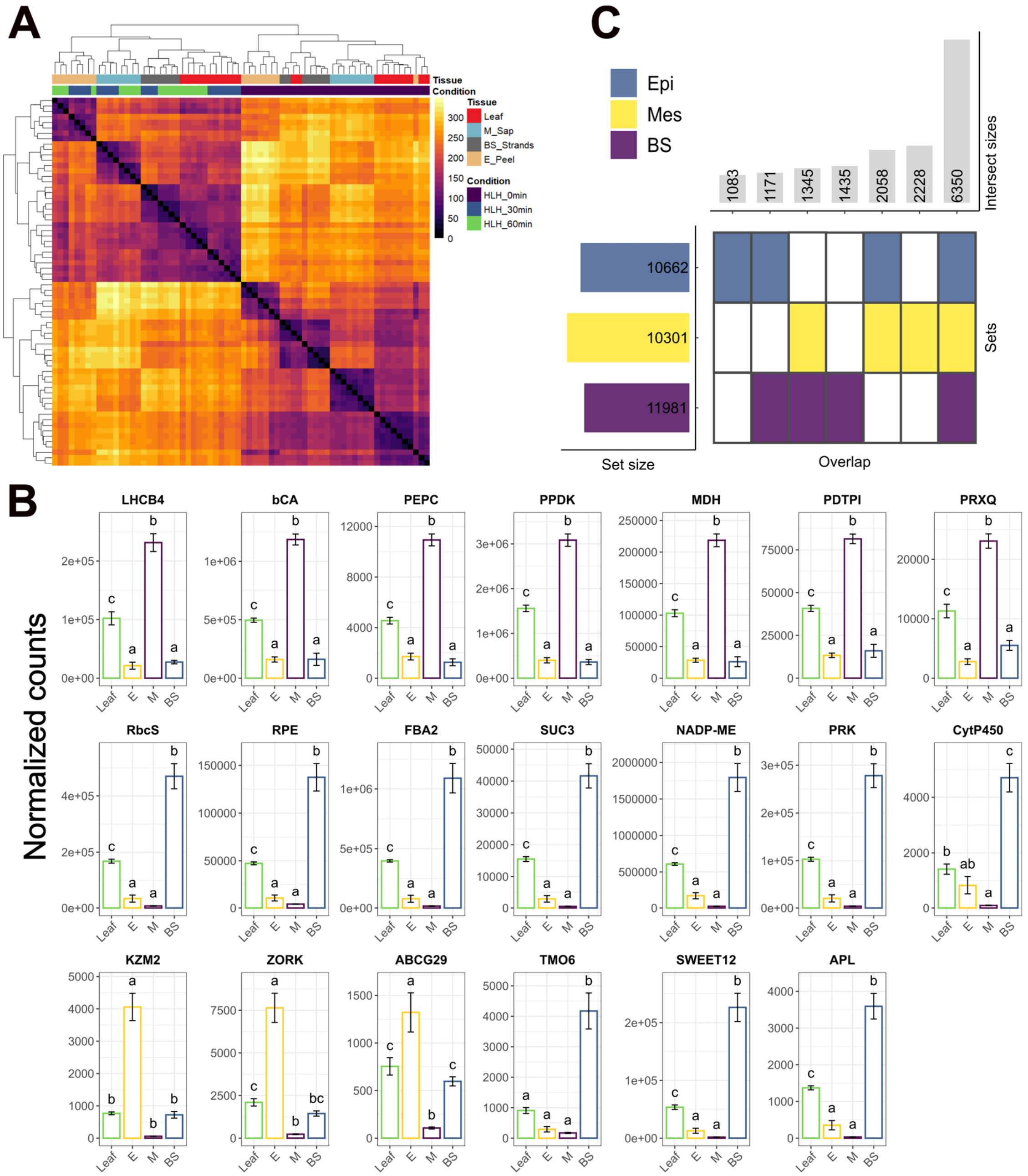


**Supplementary Figure 3. Transcriptomes of mesophyll, bundle sheath, and epidermal cell fractions are consistent with previously described functions.**
**(A)** Hierarchical clustering of RNA-seq biological replicates. Colour scale represents Euclidean distance between samples (lower indicating greater similarity). **(B)** RNA-seq expression data (as normalized counts) of mesophyll marker transcripts (top row; *LHCB4*, *Light-harvesting chlorophyll a/b binding 4*; *CA*, *Carbonic anhydrase*; *PEPC*, *Phosphoenolpyruvate carboxylase*; *PPDK*, *Pyruvate phosphate dikinase*; *MDH*, *Malate dehydrogenase*; *PDTPI*, *plastid triose phosphate isomerase*; *PRXQ*, *Peroxiredoxin Q*), bundle sheath marker transcripts (middle row; *RBCS*, *Rubisco small subunit*; *RPE*, *Ribulose-5-phosphate-3-epimerase*; *FBA2*, *Fructose-1,6-bisphosphate aldolase 2*; *SUC3*,  *Sucrose transporter 3*; *NADP-ME*, *NADP-malic enzyme*; *PRK*, *Phosphoribulokinase*;

*CYP78A6*), and guard cell (*KZM2, Potassium channel* Zea mays *2*), subsidiary cell (*ZORK,* Zea mays *Outward-Rectifying potassium channel*), epidermal pavement cell (*ABCG29, ATP-Binding Cassette Subfamily G member 29*), procambium (*TMO6, Target of monopteros 6*) , phloem (*SWEET12, Sugar will eventually be exported transporter 12*) and companion cell (*APL, Altered phloem development*) marker transcripts (bottom row) in whole leaf, epidermal peels (E), mesophyll sap (M), and bundle sheath strands (BS). Letters indicate statistically significant differences (One-way ANOVA followed by Tukey's HSD test, $p < 0.05$). **(C)** UpSet plot summarizing the number and overlap of DEGs between epidermal peels (E), mesophyll sap (M) and bundle sheath strands (BS).

#### Supplementary Figure 4

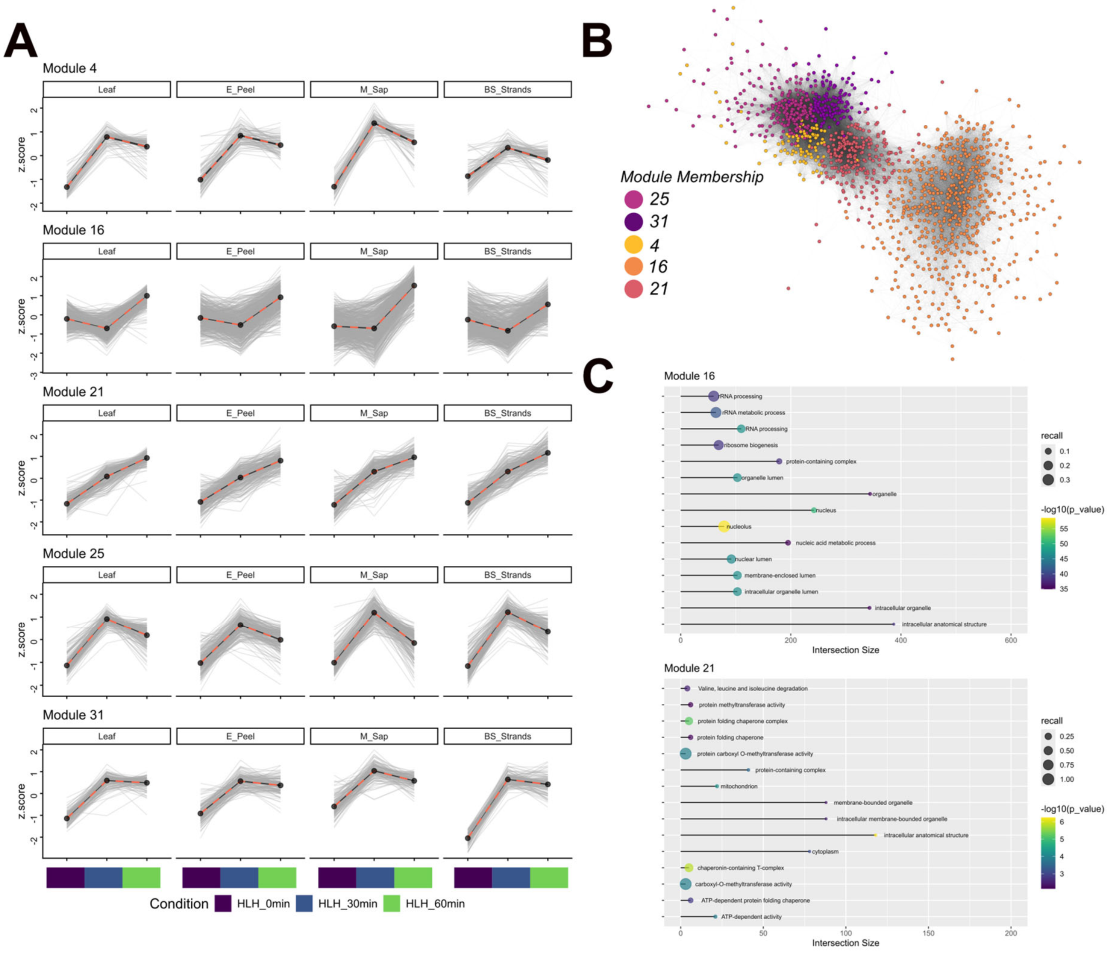


**Supplementary Figure 4. Transcripts related to ROS detoxification and chloroplast signalling are not represented within core sets of high light and heat-responsive DEGs.**
**(A)** Gene expression profiles of co-expression modules not associated with specific cell types. Grey lines represent the expression patterns of each gene within the module (visualized as the genes Z-score over the course of HL+H treatment), and red dashed lines the mean expression calculated from all genes within the module. **(B)** Co-expression subnetwork of genes from modules shown in **(A)**. Each node represents a gene (coloured by its module membership within the broader network), and edges significant correlations ($R > 0.7$) in transcript levels between two genes. Darker edges indicate higher correlations between transcripts. **(C)** The top 15 most enriched gene ontology terms in co-expression modules 16 and 21. Colour scale represents the -$Log_{10}$ transformed false discovery rate (FDR), point size the recall ratio (number of genes in dataset belonging to the GO term/total number of genes belonging to the GO term), and intersection size (the total number of input genes classified under each GO term) is plotted on the x-axis.

**Supplementary Figure 5**

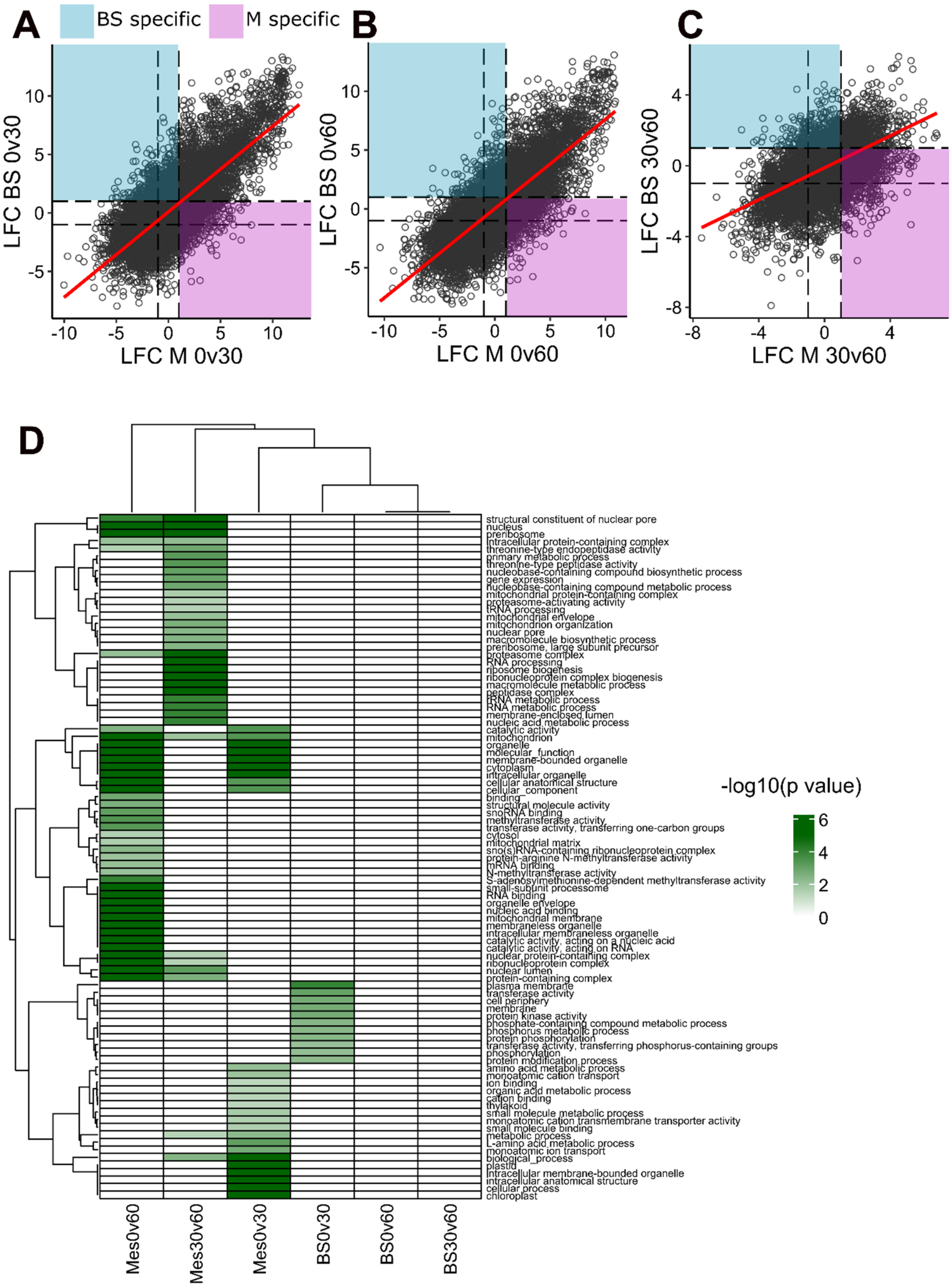


**Supplementary Figure 5. Distinct responses of mesophyll and bundle sheath cells to high light and heat stress.**

**(A-C)** Plots comparing $\log_2$ fold-change of DEGs in mesophyll and bundle sheath cells at different stress timepoints used to identify M- and BS-specific DEGs. **(D)** Heatmap of gene ontology terms enriched in cell-specific DEGs, comparing 0 vs 30, 0 vs 60, and 30 vs 60 HL+H timepoints. Colour scale represents the -$\log_{10}$ transformed *p* values corrected for false discovery rate.

#### Supplementary Figure 6

A

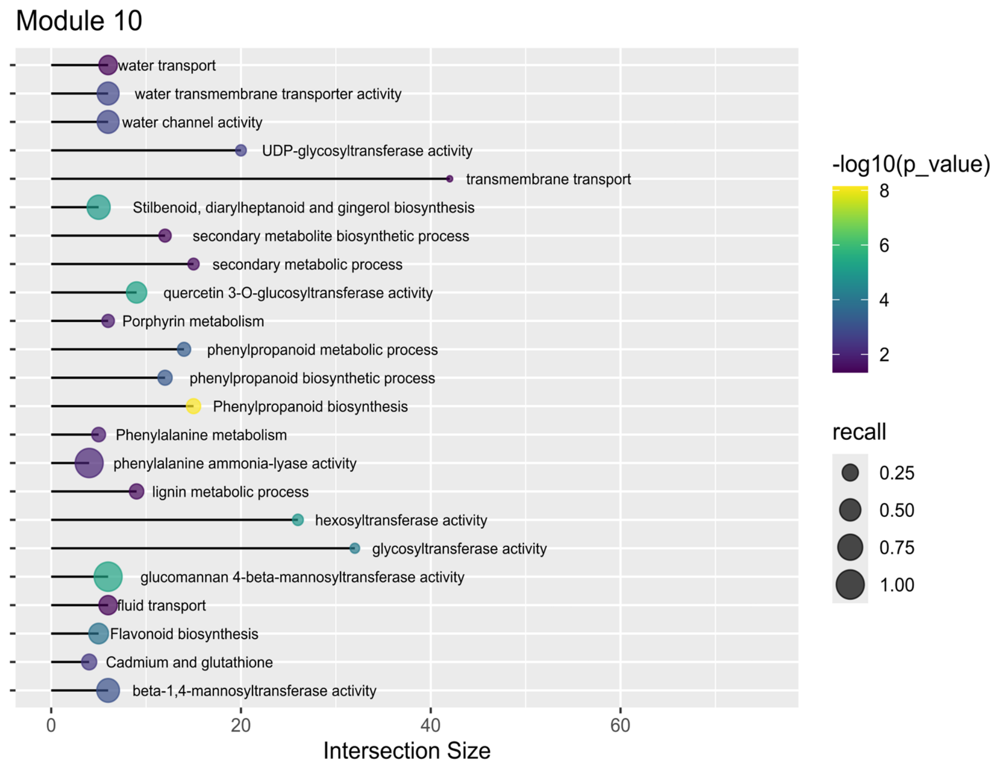


B

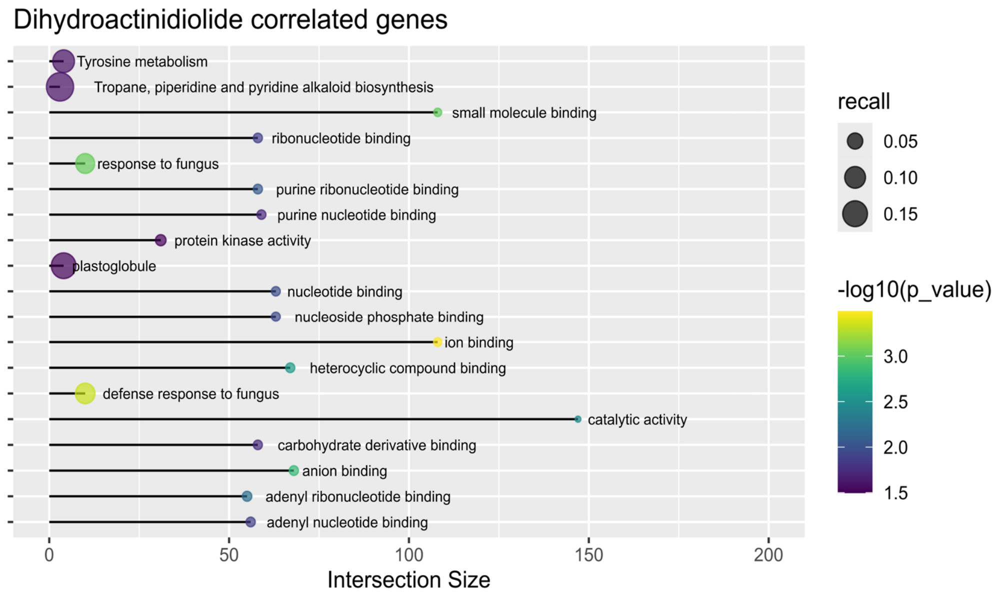


**Supplementary Figure 6. Enriched Gene Ontology (GO) terms in mesophyll-enriched and DHA-correlated gene expression changes.**
**(A)** Bubble plot of all GO terms enriched within module 10. Colour scale represents the -$Log_{10}$ transformed false discovery rate (FDR), and intersection size (the number of genes belonging to each GO term) plotted on the x-axis. **(B)** Bubble plot of all GO terms enriched in DHA-correlated genes. Colour scale represents the -

$Log_{10}$ transformed false discovery rate (FDR), point size the recall ratio (number of genes in dataset belonging to the GO term/total number of genes belonging to the GO term), and intersection size (the number of genes belonging to each GO term) plotted on the x-axis.

**Supplementary Figure 7**

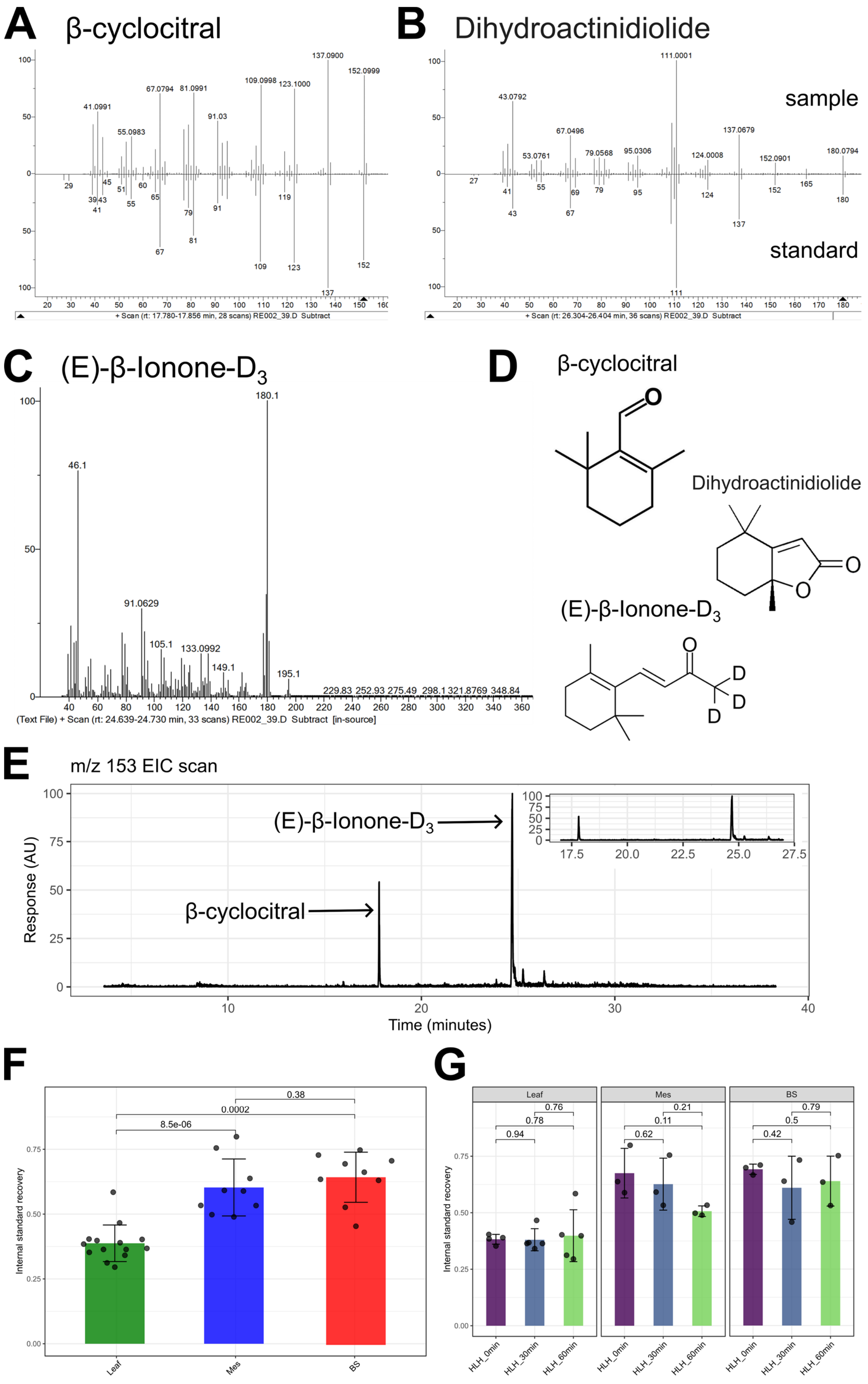

**Supplementary Figure 7. Identification of volatile apocarotenoids by HSPME-GCMS and recovery of internal standard across treatments and tissues.**

**(A-C)** Mass spectra of volatile apocarotenoids detected in plant samples and their comparison to authentic standards. **(D)** Structures of β-cyclocitral, dihdydroactinidiolide, and the $D_3$-β-ionone internal standard. **(E)** Representative scan of a mesophyll sap sample at m/z 153, the quantifier ion of β-cyclocitric acid. No peaks had spectra consistent with those previously published for β-cyclocitric acid. Inset panel shows zoomed area between β-cyclocitral and $D_3$-β-ionone peaks. **(F)** Recovery of internal standard across the 3 tissue fractions **(G)** and over the course of HL+H treatment. Internal standard recovery is expressed as the peak area of $D_3$-β-ionone spiked into plant samples compared to its peak area in a sample containing only 20% NaCl. *p* values are derived from either one- or two-way ANOVA followed by Tukey's post-hoc test ($p < 0.05$).

**Supplementary Figure 8**

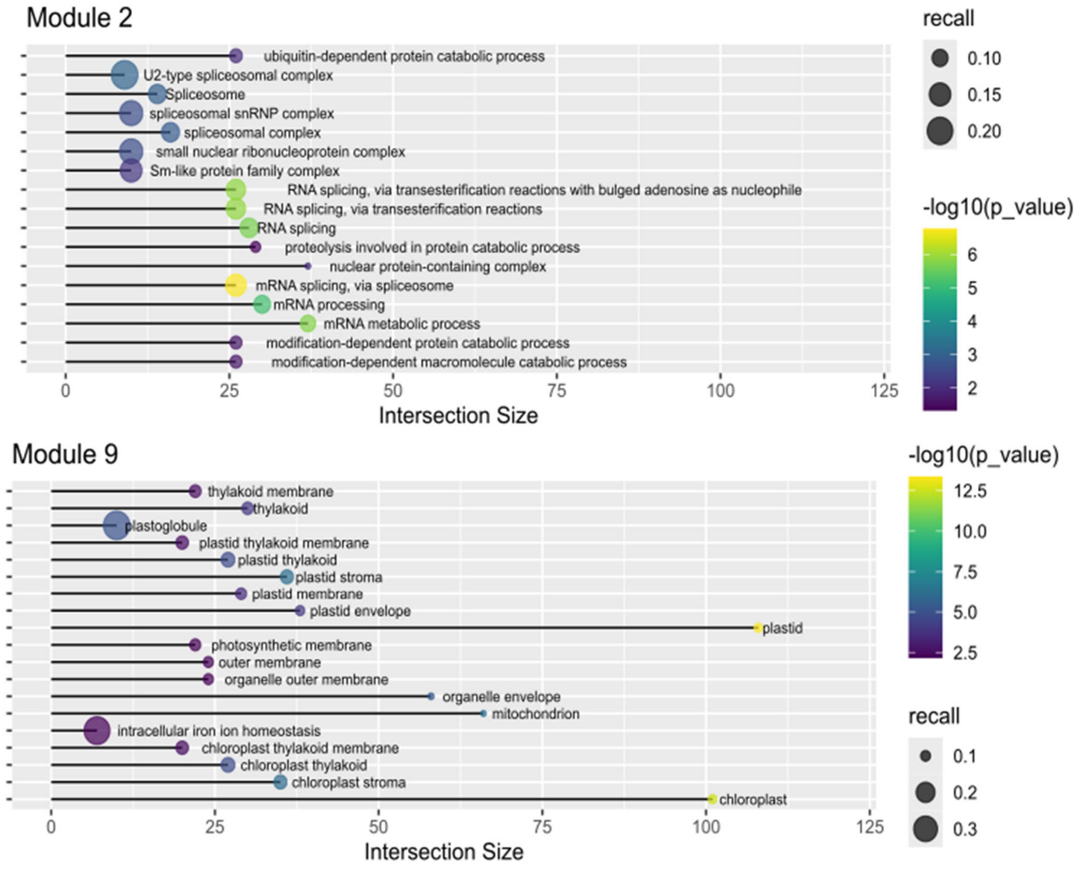


**Supplementary Figure 8. Enriched Gene Ontology (GO) terms in two mesophyll-preferential DEG modules showing significant correlation with PAP levels.** Bubble plot of all GO terms enriched within co-expression modules 2 and 9. Colour scale represents the -Log10 transformed false discovery rate (FDR), point size the recall ratio (number of genes in dataset belonging to the GO term/total number of genes belonging to the GO term), and intersection size (the number of genes belonging to each GO term) plotted on the x-axis.

**Supplementary Figure 9**

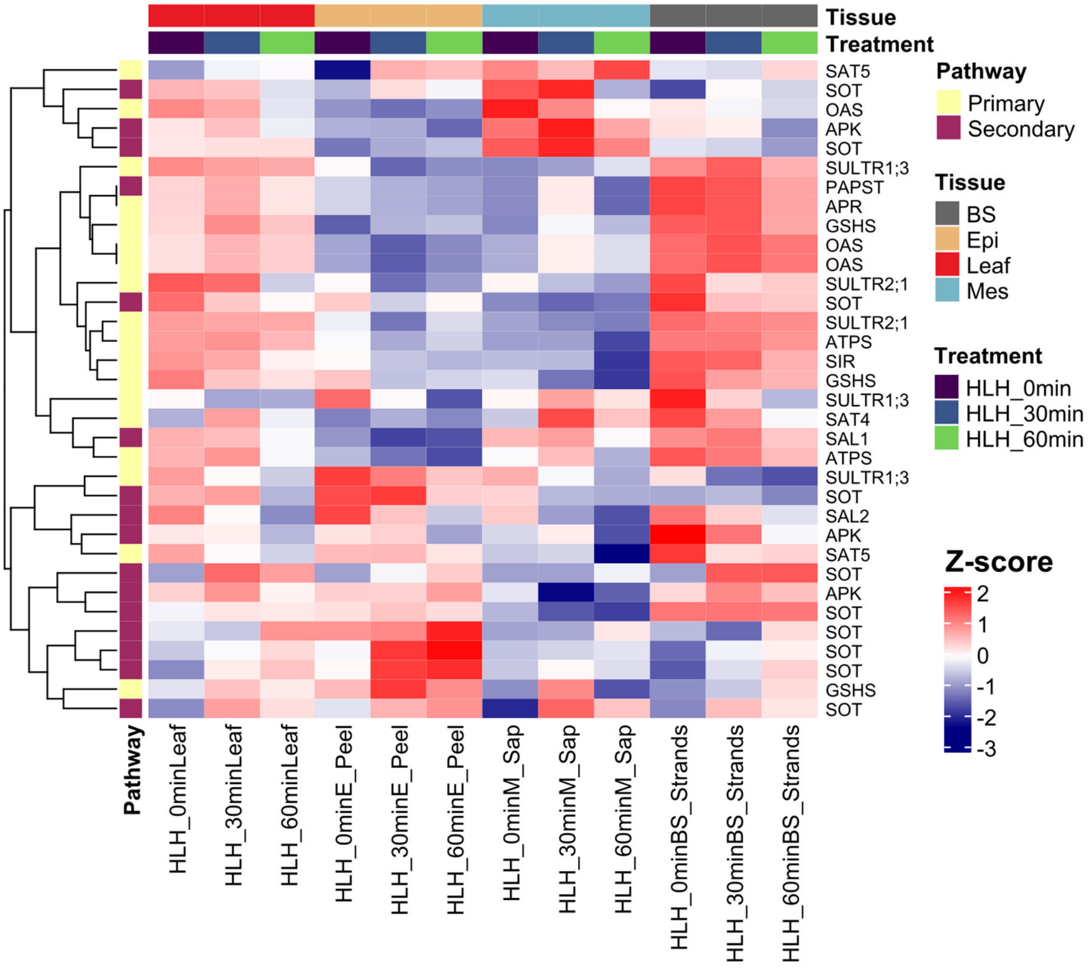


**Supplementary Figure 9. *S. viridis* exhibits differential partitioning of genes for primary and secondary sulfur metabolism and two distinct *SAL* genes.** Expression profiles of transcripts implicated in primary and secondary sulfur metabolism (represented as scaled Z-scores).

#### Supplementary Figure 10

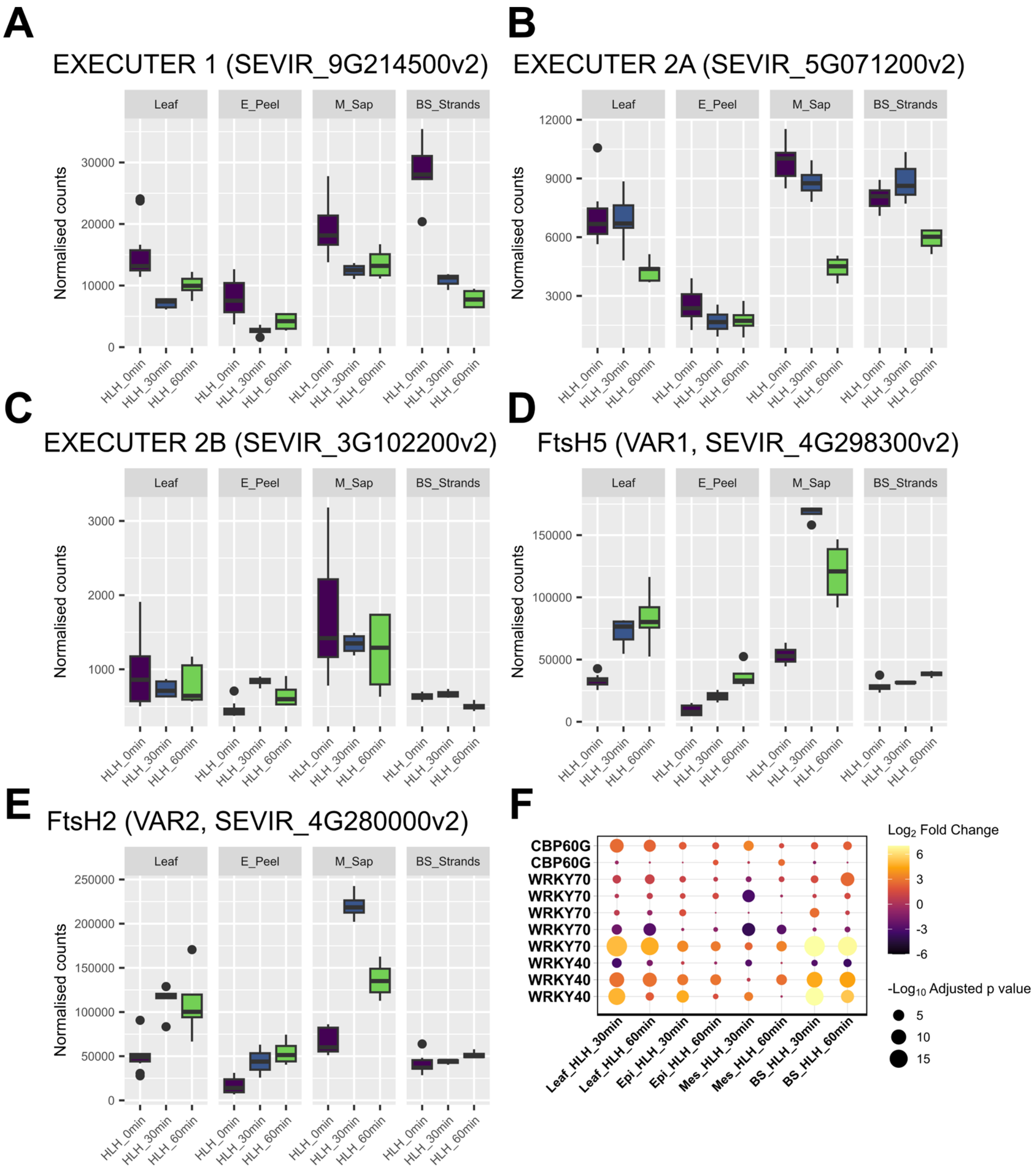


**Supplementary Figure 10. Components of EXECUTER1/2 signalling are distributed evenly between mesophyll and bundle sheath cells.**
RNA-seq expression data (as normalized counts) of the EXECUTER 1 **(A)**, two EXECUTER 2 homologs (**B-C**), FtsH5 (**D**) and FtsH2 (**E**) protease transcripts in different cell types and in response to HL+H treatment. **(E)** Expression bubble plot of genes related to EXECUTER-dependant chloroplast signalling in response to HL+H. The colour scale represents the $Log_2$-fold change in expression compared to 0 minutes HL+H treatment, and dot size the significance of this change as $-Log_{10}$ of the adjusted *p*-value.

#### Supplementary Figure 11

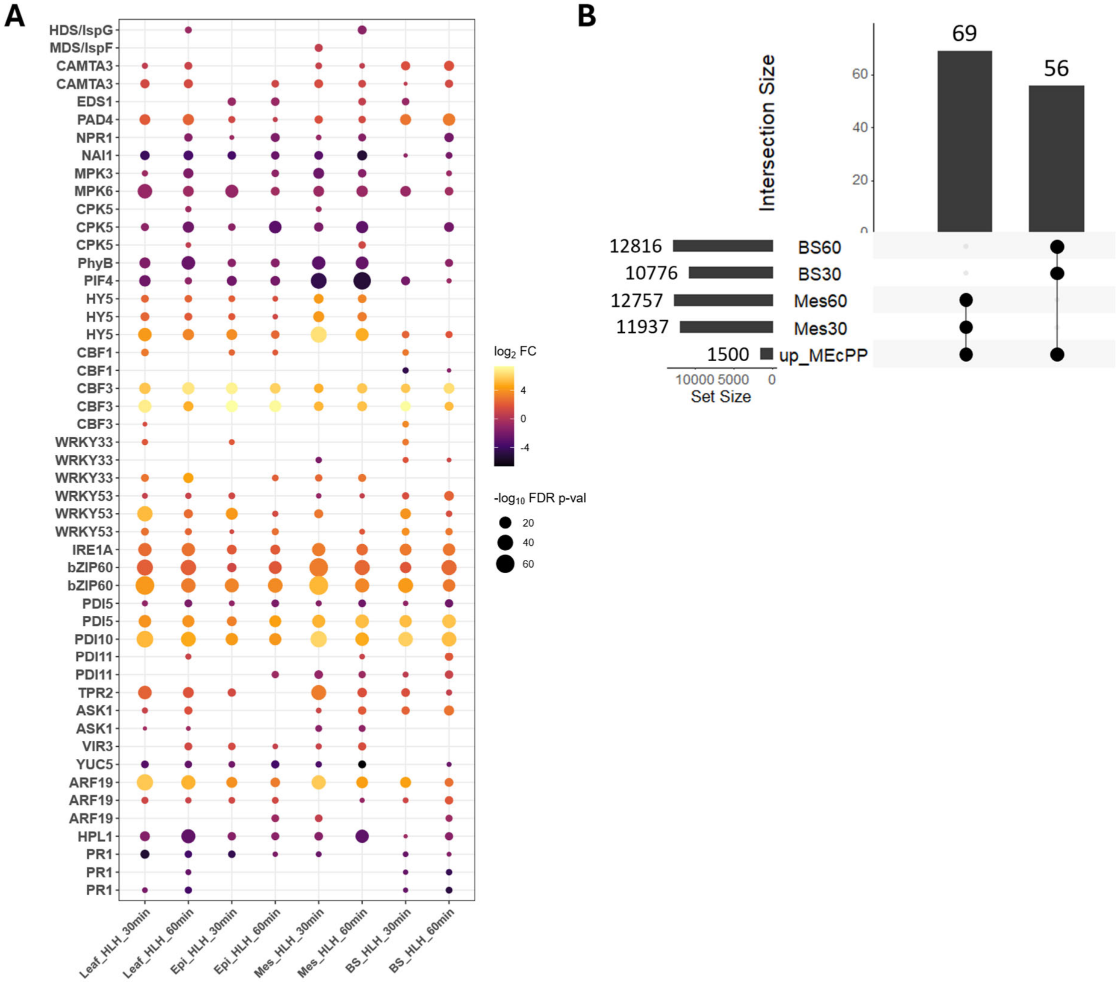


**Supplementary Figure 11. Expression of marker genes associated with MEcPP-dependent chloroplast signalling. (A)** Expression bubble plot of MEcPP-responsive genes in response to HL+H treatment. The colour scale represents the $Log_2$-fold change in expression compared to 0 minutes HL+H treatment, and dot size the significance of this change as $-Log_{10}$ of the adjusted *p*-value. **(B)** UpSet plot illustrating distinct intersects between DEGs enriched in *S. viridis* Mes sap or BS strands after 30 or 60 minutes of HL+H treatment compared to the MEcPP-responsive DEGs identified in the Arabidopsis *ceh1* mutant by Bjornson et al (2017).

**Supplementary Figure 12**

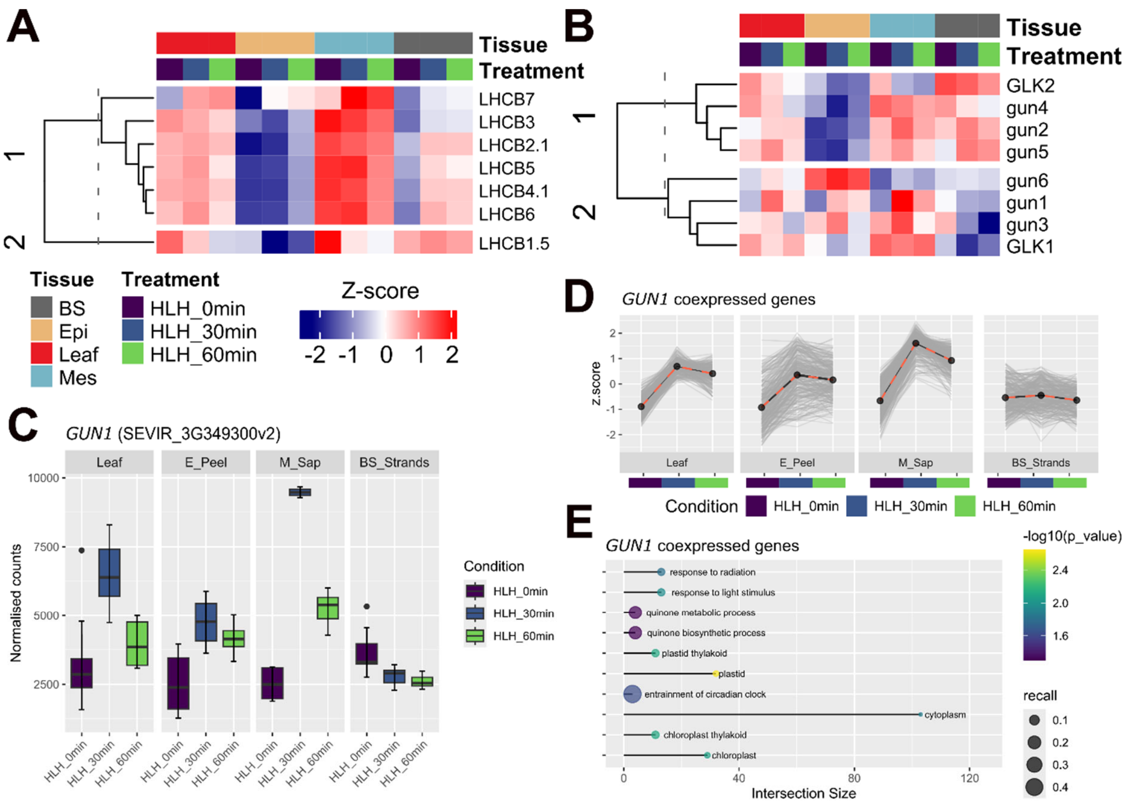


**Supplementary Figure 12. Both mesophyll and bundle sheath cells have capacity for *GUN*-mediated chloroplast signalling.** Clustered heatmaps showing Z-scaled expression values of light harvesting complex B (*LHCB*) components (**A**) and *GUN*/*GLK* transcripts (**B**) across tissues and treatments. (**C**) RNA-seq expression data (as normalized counts) of the *S. viridis GUN1* gene. (**D**) Expression profiles of *GUN1* co-expressed transcripts. Grey lines represent the expression patterns of each gene within the module (visualized as the genes z-score over the course of HL+H treatment), and red dashed lines the mean expression calculated from all genes within the module. (**E**) All GO terms enriched in this gene set. Colour scale represents the -$Log_{10}$ transformed false discovery rate (FDR), point size the recall ratio (number of genes in dataset belonging to the GO term/total number of genes belonging to the GO term), and intersection size (the total number of input genes belonging to each GO term) is plotted on the x-axis.

#### Supplementary Figure 13

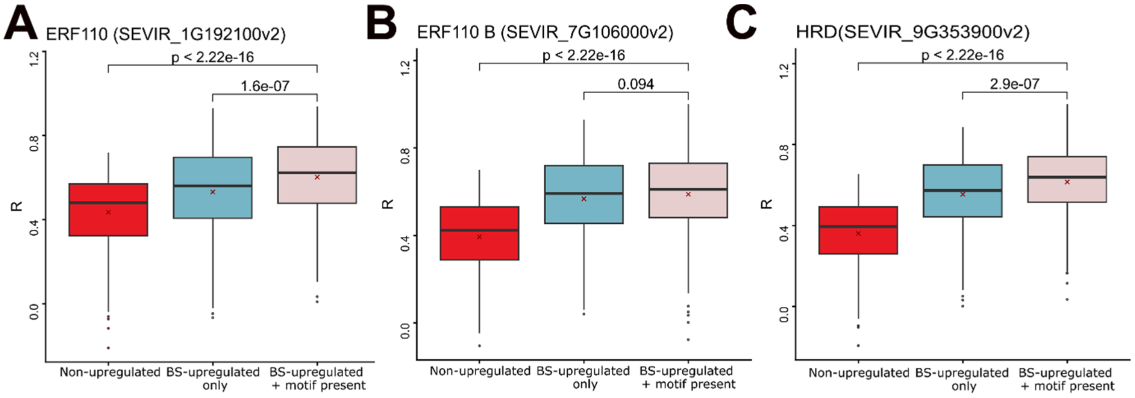


**Supplementary Figure 13.** Additional pairwise correlation plots between the transcript levels of **(A-B)** ERF110 and **(C)** HRD homologs and their putative targets within co-expression module 3. The correlation between AP2/ERF and downstream target gene expression was compared between 3 groups: transcripts that contained the binding motif in **Fig 4C** in their upstream promoter regions (red boxes), other BS-upregulated transcripts from modules 3 and 20 that did not contain the motif in their promoters (blue boxes), and randomly selected transcripts from outside of module 3 (pink boxes) ($N$ = 438 for each group). Statistical significance between groups was determined via Student's two-tailed $t$-test ($p < 0.05$) and corrected for multiple comparisons using the Benjamini-Hochberg procedure.

#### Supplementary Figure 14

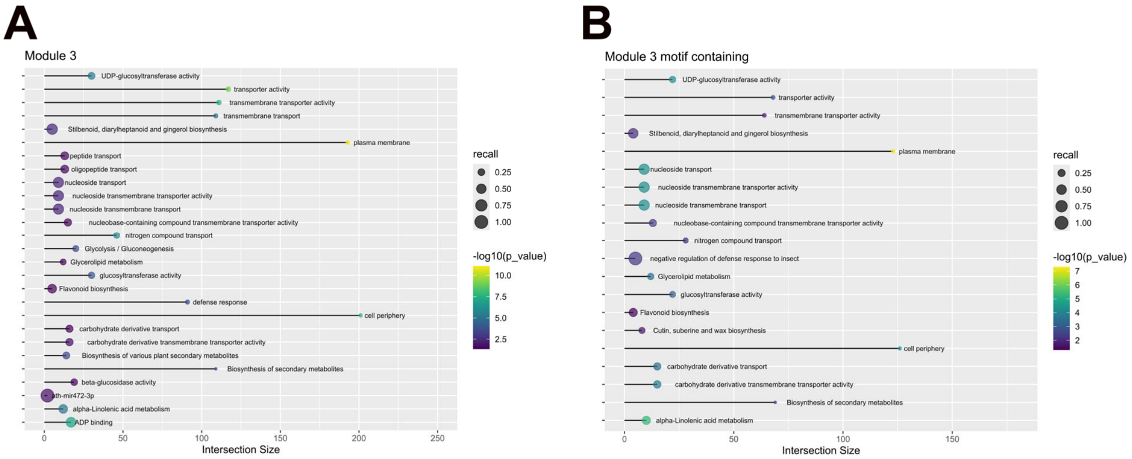


**Supplementary Figure 14. Bundle sheath-upregulated transcripts are enriched in carbohydrate and secondary metabolism.**
All enriched gene ontology terms within co-expression module 3 **(A)**, and the subset of co-expression module 3 transcripts containing the AP2/ERF binding motif in their upstream promoter regions **(B)**. Colour scale represents the -$\text{Log}_{10}$ transformed false discovery rate (FDR), point size the recall ratio (number of genes in dataset belonging to the GO term/total number of genes belonging to the GO term), and intersection size (the total number of input genes belonging to each GO term) is plotted on the x-axis.

**Supplementary Figure 15**

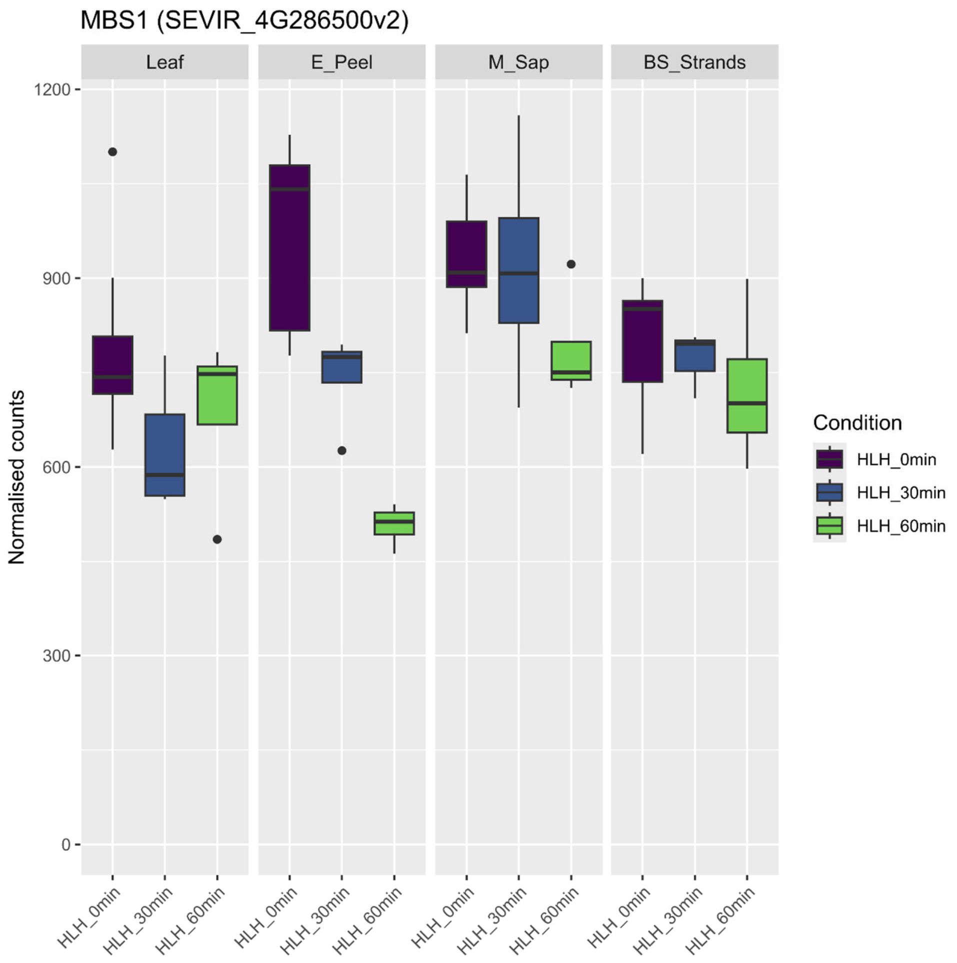


**Supplementary Figure 15. Expression of *MBS1* transcript across cell types and stress treatments.**

RNA-seq expression data (as normalized counts) for MBS1 transcripts detected in different cell types and in response to HL+H treatment. Box and whisker plots are represented as the median with upper and lower quartiles, while dots represent outlier counts.

#### Supplementary Figure 16

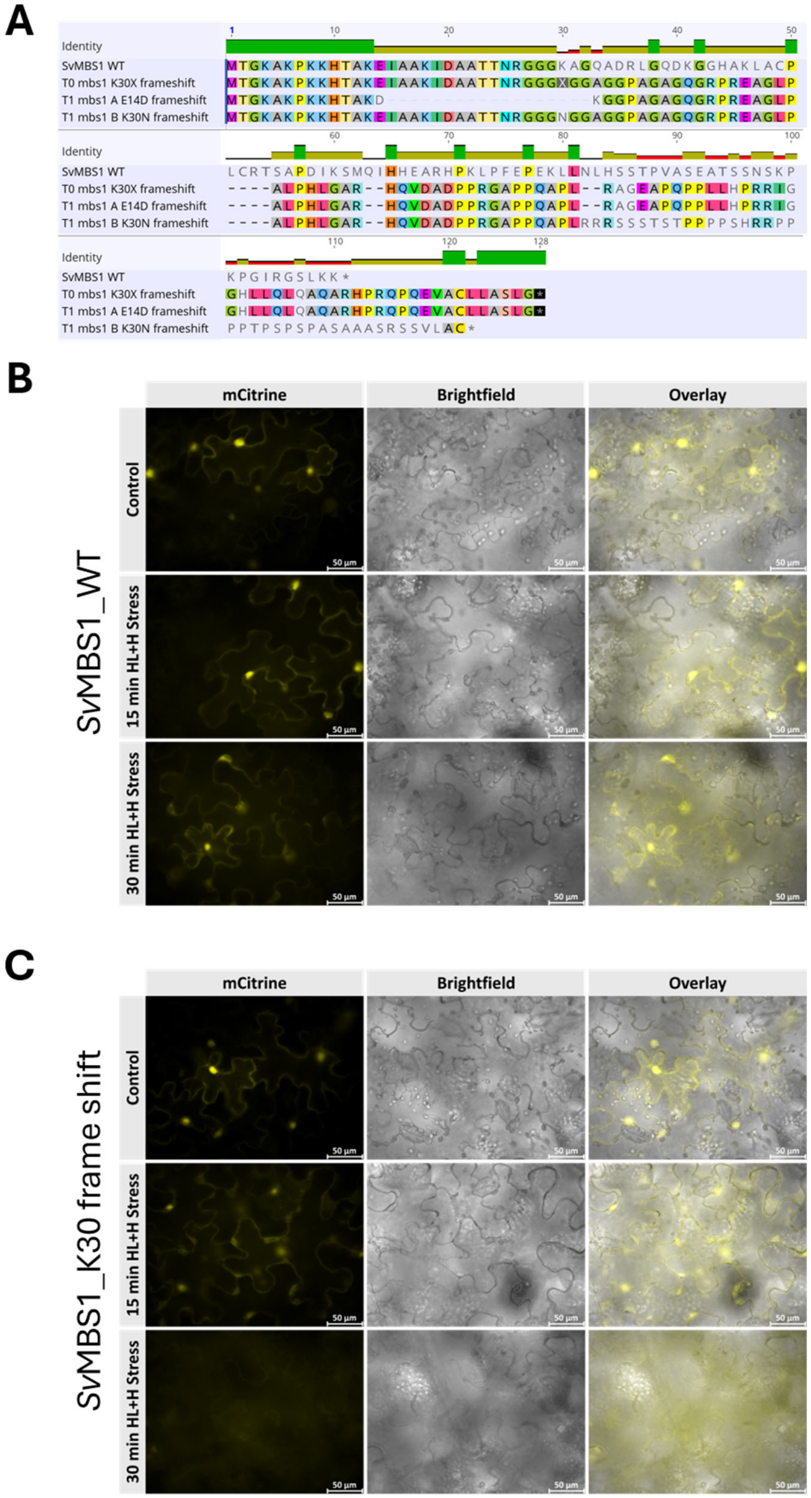


**Supplementary Figure 16. Effect of MBS1 mutations on protein abundance during HL+H treatment** **(A)** Amino acid alignment of identified CRISPR-Cas9 gene-edited *MBS1* mutants to WT in *S. viridis*. The K30X line was the parental ($T_0$) line from which the E14D (*mbs1 A*) and K30N (*mbs1 B*) mutants were identified in the subsequent $T_1$ generation. The E14D and K30N mutants were used in Fig 5. **(B)** WT MBS1-mCitrine protein fusions transiently expressed in *N. benthamiana* leaves, which were then exposed to control or

HL+H treatment. MBS1-mCitrine fluorescence is false-coloured yellow. Scale bars represent 50 µm. (**C)** MBS1 K30X frame shift mutated protein transiently expressed in *N. benthamiana* leaves, which were then exposed to control or HL+H treatment. MBS1-mCitrine fluorescence is false-coloured yellow. Scale bars represent 50 µm.

**Table S1: Oligonucleiotide sequences designed to target *MBS1*, determine edits in *MBS1* and amplify *mbs1* null mutant alleles.**

NGG PAM sequences downstream of guides for SpCas9 recognition and binding shown in bold.

| **Purpose** | **Oligonucleotide** | **Sequence (5' - 3')** |
|---|---|---|
| CRISPR/Cas9 editing of *MBS1* | SvMBS1 gRNA 1 | AACCGTGGCGGCGGGAAGGC **GGG** |
| | SvMBS1 gRNA 2 | AGGAGCTTCTCCGGCTCGAA **GGG** |
| Screening edited *mbs1* lines | SvMBS1 F | CATCCTCTCGCACATCACCGC |
| | SvMBS1 R | CGATGGATTAATTATAGATCGGGTTGGTGGATAG |
| Amplifying *mbs1* null alleles for transient expression | SvMBS1 WT CDS F | AAGAAGACAACTCAAATGACGGGCAAGGCGAA |
| | SvMBS1 A CDS R | AAGAAGACAACTCGCGAACCCAAGCTAGCAAGCAAGC |
| | SvMBS1 B CDS R | AAGAAGACAACTCGCGAACCGCAAGCAAGCACGCTACTTC |
